# Mechanisms of punctuated vision in fly flight

**DOI:** 10.1101/2020.12.16.422884

**Authors:** Benjamin Cellini, Wael Salem, Jean-Michel Mongeau

## Abstract

To guide locomotion, animals control their gaze via movements of their eyes, head, and/or body, but how the nervous system controls gaze during complex motor tasks remains elusive. Notably, eye movements are constrained by anatomical limits, which requires resetting eye position. By studying tethered, flying flies (*Drosophila*) in a virtual reality flight simulator, we show that ballistic head movements (saccades) reset eye position, are stereotyped and leverage elastic recoil of the neck joint, enabling mechanically assisted redirection of gaze. Head reset saccades were of proprioceptive origin and interrupted smooth movements for as little as 50 ms, enabling punctuated, near-continuous gaze stabilization. Wing saccades were modulated by head orientation, establishing a causal link between neck signals and execution of body saccades. Furthermore, we demonstrate that head movements are gated by behavioral state. We propose a control architecture for biological and bio-inspired active vision systems with limits in sensor range of motion.

## INTRODUCTION

Animals move their eyes, head and/or body to stabilize and shift gaze. In particular, eye movements display a variety of dynamic modes, consisting of interacting slow- and fast-phases (Land, 1992). In response to continuous motion, human eyes will alternate between periods of slow stabilizing movements and rapid, anti-directional shifts in gaze called “saccades”, a motor pattern generally referred as optokinetic nystagmus (Purves et al., 2004). Periods of slow eye movements and saccades are widespread in the animal kingdom, observed in primates, birds, fish, crabs, and a variety of insects, although the physiological mechanisms enabling eye movements vary across phyla (Land, 2019). In insects, the eyes cannot move independently of the head, thus saccades are performed via both head and body movements (Collett and Land, 1975; Land, 1973). Revealing the strategies insects use to control gaze, specifically how and why they perform saccades, can provide unique insights into how eye movements are coordinated in locomoting animals.

Analogous to humans, in flying flies a rotating cylinder with vertical stripe elicits smooth movement and rapid reset saccades of the body and head (Collett and Land, 1975; Land, 1973; Mongeau and Frye, 2017). Similarly, as in humans, fly vision is also punctuated, combining continuous and discrete movements (Cellini and Mongeau, 2020a; Collett and Land, 1975; Land, 1973; Mongeau et al., 2019; Mongeau and Frye, 2017). Functionally, reset saccades prevent the eyes from reaching their anatomical limits, which is a fundamental constraint on gaze stabilizing systems across phyla. Healthy humans can move their eyes ~±45° about yaw (Shin et al., 2016), whereas in *Drosophila* head movements about yaw have a limit of about ~±15° from body midline (Cellini and Mongeau, 2020b; Fox and Frye, 2014). A visual control system combining smooth movement and reset saccades could therefore overcome these classes of strict anatomical limits. Whereas the interaction between smooth body movements and body saccades in flies has been revealed, how head movements contribute to gaze stabilization remains poorly understood. Head saccades are generally coupled with body saccades in flies and in bees and may aid in reducing motion blur during a body saccade (Boeddeker et al., 2010; Schilstra and van Hateren, 1998), however the dynamics and control of head saccades during gaze stabilization remain elusive. How are head smooth movements and saccades coordinated and how are head reset saccades triggered? To what extent are head and wing saccades coupled? Revealing these interactions are critical to understanding the punctuated nature of fly vision and coordination in flight, and ultimately its underlying neural control circuitry.

We investigated saccadic head and wing movements in rigidly tethered *Drosophila* during gaze stabilization. Because the head and wings displayed distinct smooth and saccadic dynamics we modeled the visuomotor response as a hybrid (switching) system (Cellini and Mongeau, 2020b; Goebel et al., 2012). We confirm that during continuous visual motion, a stereotyped reset saccade is triggered when the head nears its anatomical limit, wherein the head moves in the opposite direction of the visual stimulus. Flies leveraged passive elastic forces of the neck joint, providing an instantaneous acceleration at the start of a head reset saccade. In between saccades, flies reduced retinal slip by up to 70% via smooth head movements. This stabilize-and-reset strategy enabled flies to stabilize gaze quasi-continuously, overcoming neck joint anatomical limits. By correlating saccade rate with the head’s excursion, we uncovered the role of proprioceptive feedback from the neck sensory system in generating saccades. Head and wing saccades were triggered concurrently but fixing the head reduced the wing saccade rate, further implicating proprioceptive feedback. Finally, we demonstrate that head movements are gated by behavioral state. We outline a framework detailing how the stabilize-and-reset strategy may apply to any noncontinuous active vision system, such as mammal eyes. Altogether, our results reveal the function and mechanism of head reset saccades in an active vision system.

## RESULTS

### Head reset saccades are stereotyped and punctuate smooth movement stabilization

Free flight body saccades in flies appear stereotyped (Muijres et al., 2015; Tammero and Dickinson, 2002a). We hypothesized that head saccades would be similarly stereotyped because head and body movements are generally coordinated (Van Hateren et al., 1999). To test this hypothesis, we elicited an optomotor response in rigidly tethered flies by presenting constant velocity visual stimuli (position ramp), which has been shown to trigger head reset saccades (Figure 1A, Video 1) (Duistermars et al., 2012; Land, 1973). We developed a method to detect head saccades based on the dynamics of head position and velocity and culled saccades across experimental trials (Figure Supplement 1) (see Methods). Head saccades were interspersed between bouts of smooth movements and had a significant influence on head trajectories (Figure 1B).

**Figure 1.**
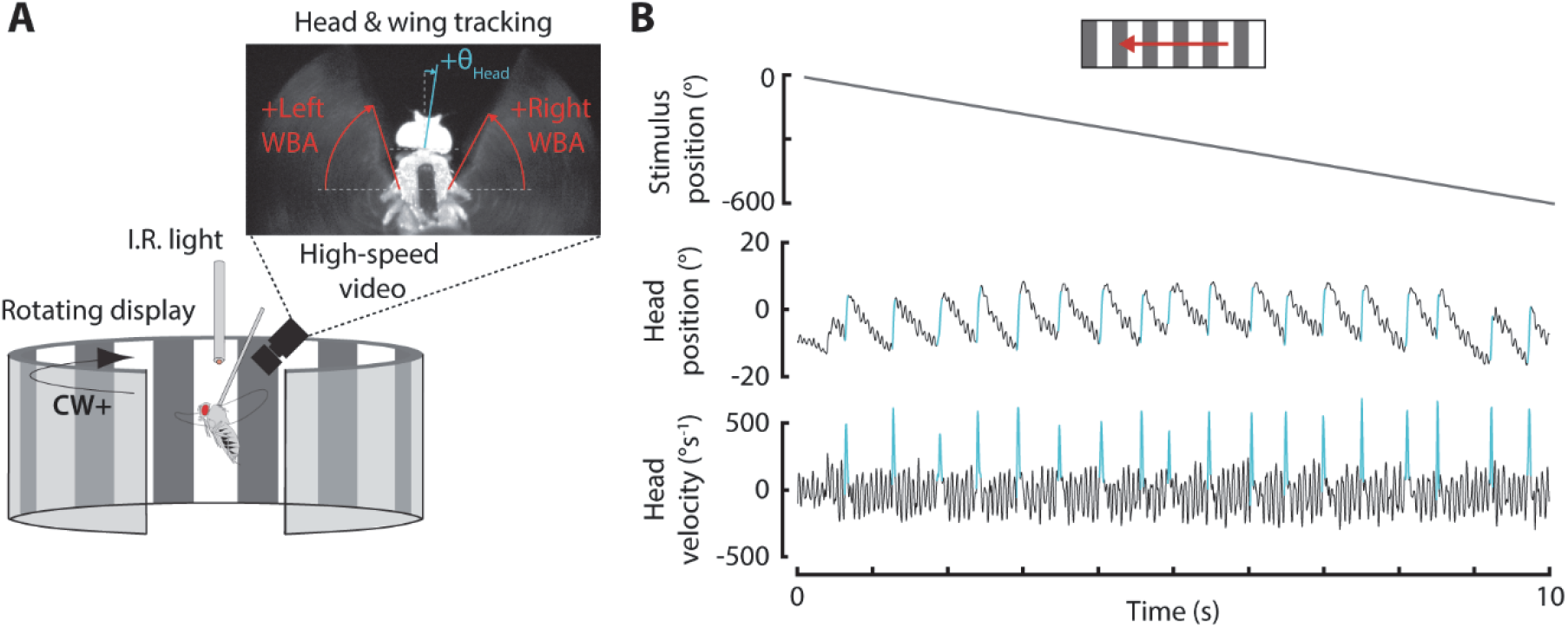
Experimental paradigm. **A)** Virtual reality flight arena. Flies were rigidly tethered to a tungsten pin and placed in the center of the arena. Head and wing movements were measured from frames acquired with a high-speed camera (200 fps) placed above the fly. The video frame shows the head orientation (cyan line) and left and right wingbeat amplitudes (red lines). **B)** An example experimental trial for a 30° spatial wavelength panorama moving at −60°s^−1^ (see Video 1). The head orientation and velocity are shown, and each saccade is highlighted.

We examined saccade dynamics by extracting each saccade and aligning it to the time it reached peak velocity (Figure 2A). The angular position and velocity of head saccades were strongly stereotyped across all stimulus speeds, consisting of a near-symmetric velocity spike lasting ~50 ms (Figure 2A). By examining the duration, amplitude, and peak velocity of saccades, we found that none of the visually elicited head saccade dynamics were robustly influenced by stimulus speed (Figure 2B). Overall, there was more variation in saccade rate. Specifically, flies performed more saccades with increasing stimulus speed up to 120°s^−1^, at which point saccade rate began to decrease. The difference in saccade rate across stimulus speeds suggests that the saccade trigger mechanism may be dependent on visual motion. Based on these findings, we conclude that head reset saccade dynamics are stereotyped but visual motion may influence their trigger.

**Figure 2.**
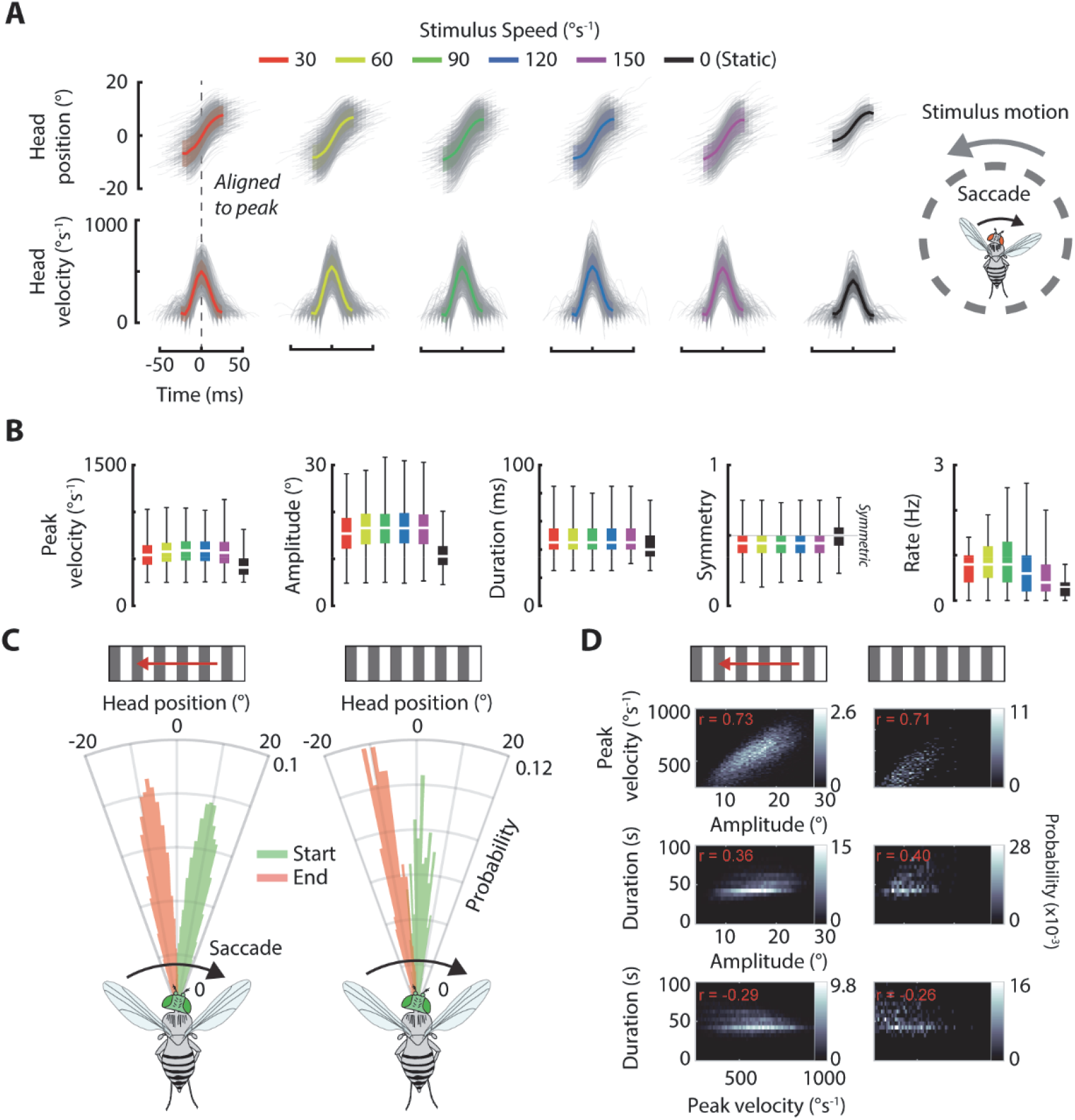
Head reset saccades are strongly stereotyped and anti-directional with respect to visual motion. **A)** Angular position and velocity trajectories of head saccades across different stimulus speeds at 30° spatial wavelength. Saccades were aligned to the peak velocity. Counter-clockwise intervals were inverted and pooled with clockwise intervals. Solid lines: mean. Shaded regions: ±1 standard deviation. Grey lines: individual saccades. **B)** Comparison of saccade dynamics and rates across stimulus speeds for 22.5°, 30°, and 60° spatial wavelengths pooled. Symmetry is the ratio of time-to-peak velocity and saccade duration, so values below 0.5 indicate the saccades start faster than they end. **C)** Polar histogram of head angular position at the start (green) and end (red) of each saccade. **D)** Relationship between saccade dynamics. *r* is the correlation coefficient. Counter-clockwise intervals were inverted and pooled with clockwise intervals. Moving stimulus: *n* = 40 flies, *N* = 15,295 saccades. Static stimulus: *n* = 8 flies, *N* = 359 saccades.

### Head saccades are anti-directional with respect to visual motion

Head saccades were triggered predominantly in the direction opposite of visual motion (Figure 2A,C). This anti-directional characteristic has been observed in previous work studying the head optomotor response in flies, locusts, and other invertebrates (Duistermars et al., 2012; Land, 2019, 1973; Shepheard, 1974). Anti-directional head saccades appear akin to “return” eye saccades in mammals that underlie optokinetic nystagmus, wherein the eyes will smoothly move in one direction before rapidly returning to the center position (Purves et al., 2004). Here we refer to anti-directional head saccades in flies as “reset” saccades, to emphasize the beginning of a new smooth gaze stabilization period after each saccade. Head reset saccades were of opposite direction to body saccades measured in a magnetic tether in previous work, wherein co-directional saccades accounted for 70% of all saccades (Mongeau and Frye, 2017), although this behavior could be influenced by restricting body motion in the rigid tether. Surprisingly, co-directional head saccades were exceedingly rare, making up less than 2% of all saccades (Figure Supplement 2). Previous work showed that anti-directional body saccade dynamics are less sensitive to visual motion speed than their co-directional counterparts, which is consistent with the head saccade tuning described here (Mongeau and Frye, 2017).

### Head reset saccades are triggered when the head is oriented near its anatomical limit

Another metric to characterize how head saccades may be initiated is the angular orientation of the head relative to the body midline. Unlike the body, which is unconstrained and able to move freely through space (unless tethered), in *Drosophila* the head has hard anatomical constraints that limit its range of motion to ~±15° about yaw (Cellini and Mongeau, 2020b; Fox and Frye, 2014). Therefore, the angular orientation of the head could play an important role in the initiation of head saccades. Head reset saccades were triggered when the head was oriented near its anatomical limit, i.e. head saccades triggered during counterclockwise rotation initiated near the left-side anatomical limit and vice-versa (Figure 2C). On average, flies engaged in smooth gaze stabilization until the head reached ±9° from body midline, then triggered a reset saccade (Figure 2C, Video 1). Head orientation at the end of a saccade was similarly clustered near the opposing side (~7.5°), moving the head through nearly its full range of motion. This is notably distinct from return eye saccade in primates, which end near the visual center (Purves et al., 2004). Taken together, these results demonstrate that head reset saccades are triggered when the head nears its anatomical limits. However, the exact sensory trigger mechanism, whether of visual or proprioceptive origin, is unclear.

### Spontaneous head saccades are distinct from visually elicited head reset saccades

We hypothesized that head saccades performed in the absence of visual motion might display different characteristics than visually elicited head saccades, as the role of the head is task dependent about the yaw axis (Fox and Frye, 2014; Geiger and Poggio, 1977). We presented flies with a static panorama and compared the dynamics of spontaneous and visually elicited reset saccades (Figure 2A–D). Compared to visually elicited saccades, the peak velocity, amplitude, and duration of spontaneous head saccades were reduced overall and were slightly more symmetric due to the similar pre- and post-peak durations (ANOVA, DOF = 5, *p* < 0.001) (Figure 2A,B). The reduced spontaneous head saccade dynamics were likely due to the more central trigger position during saccade initiation (Figure 2C). Because flies were not stabilizing a moving panorama, the head maintained a more neutral orientation. Therefore, the head was limited to a smaller range of motion amplitude when performing a saccade. For spontaneous and visually elicited saccades, amplitude and peak velocity were highly correlated (r ≈ 0.7 for both,*p* < 0.001) therefore it follows that the peak velocity of spontaneous head saccades was also reduced compared to visually elicited saccades (Figure 2D). Amplitude and duration were correlated to a lesser extent (r = 0.4, *p* < 0.001) while peak velocity and duration exhibited a slight negative correlation (r = −0.3, *p* < 0.001). Spontaneous saccades occurred at a rate more than two times lower than visually elicited saccades, demonstrating that visual motion influences the trigger of saccades (Figure 2B). Taken together, spontaneous head saccades are distinct from visually elicited saccades and appear to have a different trigger mechanism.

### Head roll during head reset saccades is feedforward

In addition to head turns in yaw, flies also rapidly rolled their head during reset saccades (Figure Supplement 4A, Video 2). Prior work demonstrated that compensatory head yaw movements are present during body mechanical roll perturbations, suggesting that yaw and roll coupling may be inherent to head control (Nalbach and Hengstenberg, 1994). The body banks during free flight saccades, thus flies simultaneously roll their head to compensate for body roll and maintain a level gaze (Schilstra and van Hateren, 1998; Van Hateren et al., 1999). Because head roll is present in rigidly tethered flies—where body movement is abolished—head roll during a saccade may consist largely of a feedforward (open-loop) motor command (Figure Supplement 2B,C). It was previously thought that compensatory head roll is reflexive, initiated primarily via feedback from flies’ mechanosensory organs—e.g. the halteres or prosternal organs—however our findings suggest that this may not be the case during saccades (Hengstenberg, 1988; National et al., 1986; Van Hateren et al., 1999). Magnetically tethered flies also roll their head during a body saccade, although body motion is restricted to yaw; however, minute body roll movement or feedback from the body yaw axis could have triggered head roll via mechanosensory feedback (Kim et al., 2017). Our results provide strong evidence that head roll is part of a feedforward motor command that could anticipate body roll during a saccade.

### Flies leverage passive elasticity of the neck joint to initiate head reset saccades

In free and magnetically tethered flight, to terminate a body saccade a fly actively brakes (decelerates) by applying a counter torque (Bender and Dickinson, 2006; Fry et al., 2003; Mongeau and Frye, 2017; Muijres et al., 2015). The profile of a head saccade displays a similar rapid deceleration that could also be indicative of active braking (Figure 3A). However, flies could rely on passive mechanics of the neck joint to damp out head motion. Furthermore, if the neck were sufficiently elastic, flies could also take advantage of spring forces to rapidly execute head saccades. To isolate the possible role of passive mechanics in head saccades and reveal the braking mechanism (active *vs*. passive), we measured the passive response of the head in anesthetized flies. In short, we displaced the head from its neutral position (0°) to near its anatomical limit, where head saccades typically begin (Figure 2C). We then rapidly released the head, allowing us to evaluate the passive response of the head-neck system (Figure 3B, Video 3). The observed passive response began at the initial displacement and rapidly moved back towards the neutral position (0°), indicative of a natural stiffness within the neck joint (Figure 3C). Interestingly, the response decayed slowly with no overshoot, thus pointing to a significant amount of damping (Figure 3C).

**Figure 3.**
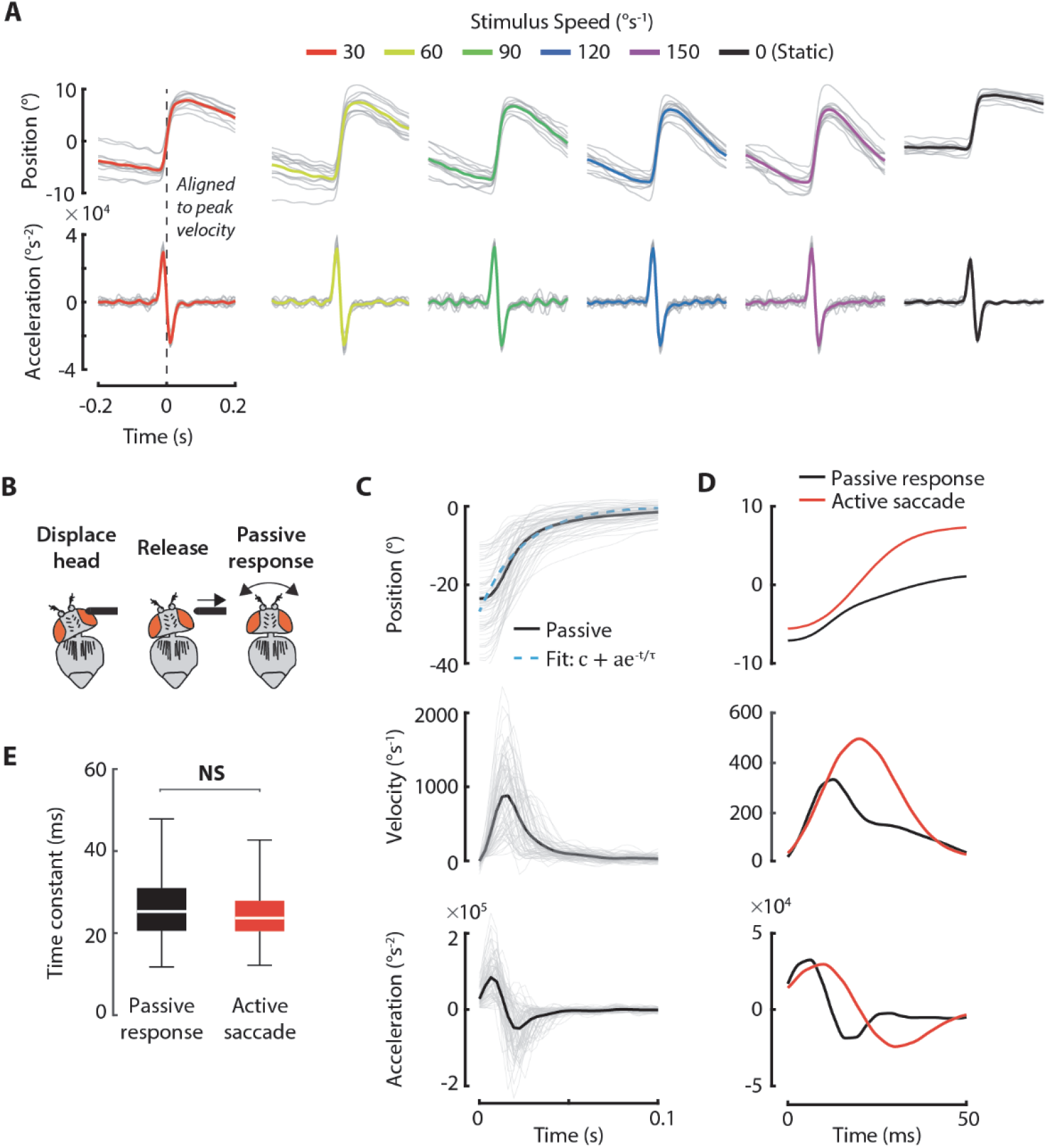
Flies leverage passive elasticity of the neck joint to initiate head reset saccades. **A)** Angular position and acceleration of head saccades across different stimulus speeds at 30° spatial wavelength. Saccades were aligned to the peak velocity. Counter-clockwise intervals were inverted and pooled with clockwise intervals. Solid colored lines: grand mean. Grey lines: individual fly means. **B)** Protocol to study the tuning of the neck joint’s passive response. The head of an anesthetized fly was displaced to near the edge of its anatomical range then subsequently released. The resulting response was then measured using custom computer vision tools. **C)** The position, velocity, and acceleration response of the head after release. Each response was fit with a 1^st^-order exponential to determine the time constant of the decay. Solid lines: mean. Grey lines: individual trials. **D)** Comparison of mean passive response (black) with mean head active saccade trajectory (red) for a 60°s^−1^ stimulus. For comparison to active saccades, only passive responses with a starting position of less than 12° were used to compute the mean (black line). **E)** Comparison of time constants for passive responses and active saccades. There was no statistically significant difference in the time constants (*t*-test, *p* = 0.3). Passive: *n* = 6 flies, *N* = 66 trials. Active: *n* = 12 flies. *N* = 3,571 trials.

To reveal the contribution of passive mechanics during head saccades, we compared the passive head response and actively controlled head saccade trajectories (Figure 3D). For comparison, we only used passive response data for initial displacements less than 12° from the neutral position to maintain consistency with the typical head reset saccade trigger position (Figure 2C). Strikingly, the passive response and active saccade trajectories were similar within the first 10–15ms, suggesting that flies may exploit the elasticity of the neck joint when initiating head saccades (Figure 3D). Indeed, at the onset the initial acceleration could be generated passively as opposed to directly by muscle activation (Figure 3D). However, once past this initial time frame, active control would be necessary to continue moving the head, as all passively generated forces would be damped out (Figure 3D). The passive mechanics accounted for ~65% of the peak velocity achieved during head saccades (Figure 3D). Interestingly, the passive and active (saccade) peak velocity scaled similarly with the initial head orientation (passive: r = −0.8, p < 0.001, active: r = − 0.89, *p* < 0.001), further implicating passive elasticity as a mechanism for the initial acceleration in saccades (Figure Supplement 3).

To reveal the role of passive breaking as flies terminate a head saccade, we approximated the damping in the system by fitting a first-order exponential to the passive response (Figure 3B). The time constant τ was ~25 ms, roughly half the duration of an active head reset saccade (Figure 3D,E, Figure 2B). The initial displacement had little effect on the time constant (r = 0.1, *p* = 0.4), typical of a linear system (Figure Supplement 3). Remarkably, we found no significant difference between the time constants of the passive response and active saccades (*t*-test, *p* = 0.3) (Figure 3E). The similar time constants suggest that flies may rely primarily on passive damping to terminate a head saccade, although there may still be an active braking component (Figure 3D,E). Taken together, the similarity in time constants between the passive response and active saccades and the fact that the passive response can account for about two thirds of active saccade velocity suggest that flies leverage passive mechanics to rapidly accelerate the head at the onset of a saccade. Furthermore, flies may take advantage of natural neck damping to terminate a head saccade, thereby simplifying control.

### Head reset saccades enable flies to reduce retinal slip by up to 70% during smooth intervals

To elucidate the functional role of head saccades during gaze stabilization and gain further insights into their trigger, we analyzed the smooth component of head movements for moving visual stimuli (Figure 4A). Flies actively compensated for visual motion via smooth head movements, as expected during the optomotor response (Duistermars et al., 2012; Land, 1973). However, the head could only stabilize gaze for less than one second before reaching its anatomical limit (Figure 4A,B, Video 1,4). The head had a stereotyped response during inter-saccade intervals: after a preceding saccade, the head would rapidly accelerate until reaching a peak velocity, then decelerate at a lesser rate until reaching near-zero velocity, signifying the head had neared its anatomical limit (Figure 4A,B). The decreasing velocity over the length of the interval could be a result of the passive mechanical properties of the neck joint, where the restoring force at the neck increases as the head orientation deviates from the neutral orientation (Figure 3C,D). By the time the head neared the anatomical limit, a reset saccade was generally triggered (Figure 4A, 2B). Interestingly, retinal slip speed (or velocity error) had little influence on saccade initiation, as the retinal slip at the start of each saccade was approximately equal to the stimulus speed due to the attenuation of stabilizing head movements (ANOVA, DOF = 4, *p* < 0.001) (Figure 4A,C). We also calculated the built-up velocity error over time, or the integrated error, and found that this too scaled with stimulus speed (ANOVA, DOF: 4, *p* < 0.001) (Figure 4C). Unlike co-directional body saccades in a magnetic tether that were triggered in response to wide-field stimulus, the integrated error was not constant across stimulus speeds, and therefore may not serve as a trigger for head reset saccades (Figure 4C) (Mongeau and Frye, 2017). These findings therefore favor a proprioceptive-based trigger determined by the orientation of the head relative to the thorax.

**Figure 4.**
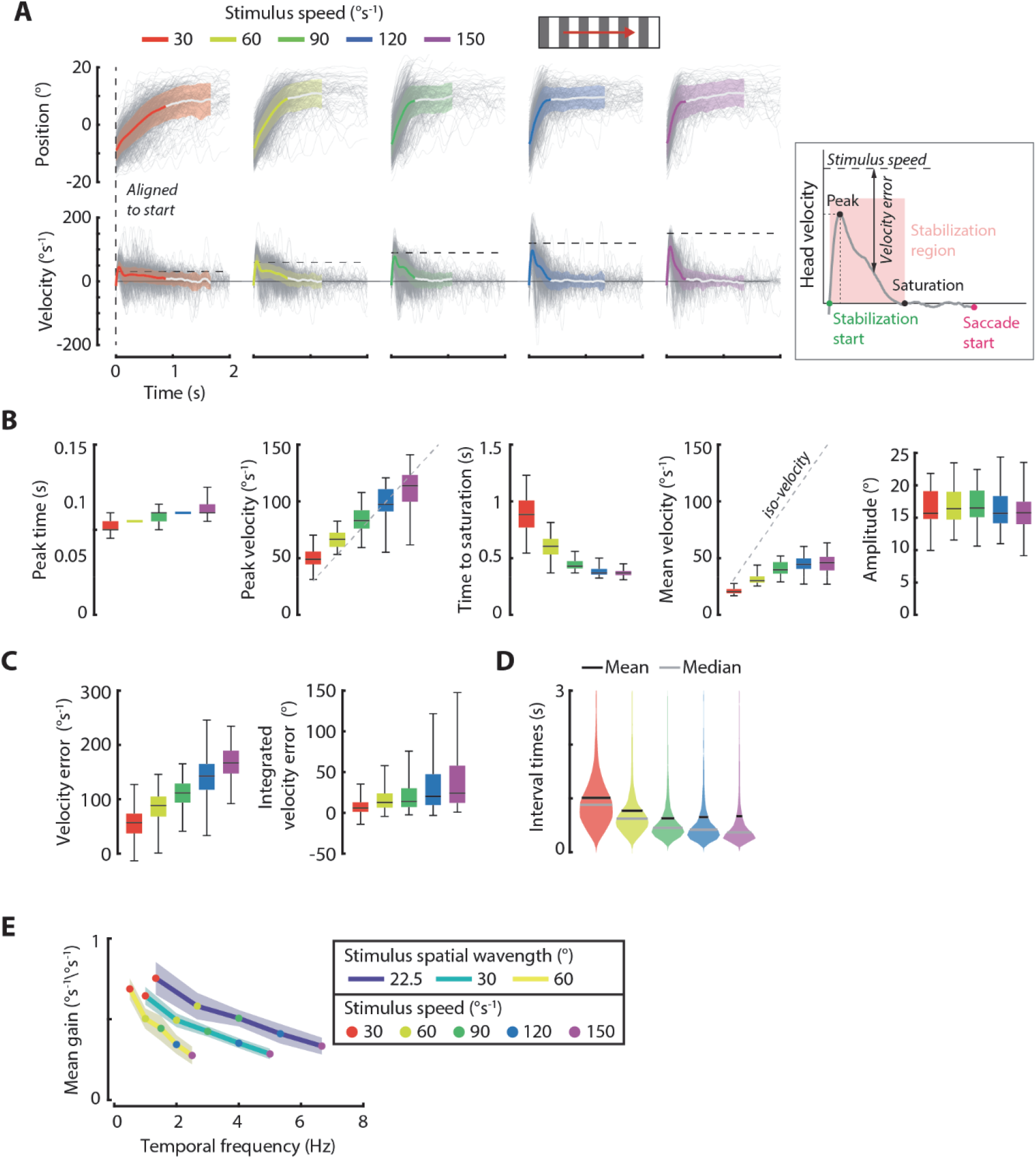
Reset saccades enable flies to reduce retinal slip by up to 70% during smooth intervals. **A)** Mean angular head position and velocity intervals across stimulus speeds for clockwise stimulus motion, at 30° spatial wavelength. Counter-clockwise intervals were inverted and pooled with clockwise intervals. Colored lines change to white after the mean interval time. Shaded regions: ±1 standard deviation. Grey lines: individual intervals. Inset: illustration of a typical smooth interval where the head rapidly accelerates, reaches peak speed, and then decelerates to near zero velocity as the head reaches its anatomical limit (saturates). **B)** Comparison of smooth head movement interval dynamics across stimulus speeds pooled across spatial wavelengths. Dynamics were extracted from the mean interval trajectory for each fly at each stimulus speed. **C)** Distributions of velocity error (retinal slip) and integrated velocity error across all intervals for each stimulus speed pooled across spatial wavelengths. **D)** Probability density estimate of smooth interval periods. We bounded the y-axis to three seconds because 98% of smooth head movement intervals fell within this range. **E)** Mean gain (mean head speed divided by stimulus speed; solid lines) for all combinations of stimulus speed and wavelength, plotted against temporal frequency. Shaded regions: ±1 standard deviation. 22.5° spatial wavelength: *n* = 16 flies, *N* = 4,409 intervals. 30° spatial wavelength: *n* = 12 flies, *N* = 3,571 intervals. 60° spatial wavelength: *n* = 12 flies, *N* = 3,817 saccades.

Head intervals were tuned to stimulus speed and head saccades enabled flies to quasi-continuously stabilize gaze. During stabilization, the initial acceleration of the head correlated with stimulus speed and reached peak speeds near the actual stimulus speed (Figure 4B). Because flies increased the speed of head movements for faster stimuli, the time the head took to reach saturation was decreased (Figure 4B). This result provides an explanation for the increasing saccade rate observed with increasing stimulus speed at lower stimulus velocities (Figure 2B), as flies would have to perform more reset saccades to continue performing stabilizing smooth movements. Without reset saccades at the end of each interval, gaze stabilization would quickly become futile, as the head would simply reach its anatomical limit and remain there. However, the hybrid, saccadic control scheme allowed flies to solve this problem by rapidly resetting their head orientation, enabling gaze stabilization to begin anew during the next smooth interval. Occasionally, the head reached its anatomical limit and remained there for a longer period (more than 1 s), thus a head saccade was not immediately triggered (Figure 4A, D). This suggests that flies may be able to inhibit the saccade reflex or that the trigger may be inherently stochastic.

To understand the extent to which flies could reduce retinal slip error during saccade-enabled smooth intervals, we computed the mean head velocity during stabilization, defined from the onset of stabilizing smooth movements to saturation (head velocity = 0 °s^−1^) (Figure 4A). We then divided the mean speed by the stimulus speed to compute a (unitless) compensatory gain (Figure 4E). On average, flies reduced retinal slip by up to 70% at low speeds and 30% at high speeds. This is in line with prior work showing that flies can compensate for oscillatory stimuli via smooth head movements by up to 60% (Cellini and Mongeau, 2020b), although the exact gain was dependent on stimulus speed and head inertial delays could have a greater influence during oscillatory motion. Compensatory head movements that reduce retinal slip have been shown to double the range of visual motion speeds encoded by the flies active vision system (head + visual system) and map the effective dynamic range of the stimulus onto the optimum of the motion vision pathway (Cellini and Mongeau, 2020b). Effectively, head movements shape and slow down inputs to the eyes. However, the effects of head movements were previously thought to be limited to stimuli moving within the head’s anatomical limits. Here we show that flies can still slow down inputs to their visual system via a combination of smooth head movements and head saccades, even for stimuli that move well beyond the anatomical limits of the head. Taken together, although reset head saccades do not directly contribute to gaze stabilization, they allow flies to stabilize gaze effectively during punctuated smooth intervals.

### Spontaneous head movements oscillate at 15 Hz in the presence of visual features

Intriguingly, in the presence of a static stimulus, flies exhibited smaller oscillatory head movements between spontaneous head saccades (Figure Supplement 5A, Video 5). These head movements contained notable frequency components between 11–18 Hz with a peak around 15 Hz, possibly part of an internal “search” strategy flies executed in the absence of visual motion (Figure Supplement 5C) (Geiger and Poggio, 1977). We measured the amplitude of these head movements to be ~2° in the 11–18 Hz frequency band across spatial wavelengths of 22.5, 30, and 60° (Figure Supplement 5D). These head movements were absent when flies were presented with a uniform background, suggesting that this search program is dependent on visual features (Figure Supplement 5B–D, Video 6)(*t*-test, *p* < 0.001). In contrast, wing movements displayed no such oscillations (Figure Supplement 5C). This is consistent with prior work showing that head and wing movements are less correlated in the absence of visual motion (Cellini and Mongeau, 2020b). The oscillatory head movements we observed are distinct from head “jitter”, which has been reported primarily in the pitch plane (Rosner et al., 2009; Van Hateren et al., 1999). Jitter frequencies in the studies were concentrated near wingbeat and haltere stroke frequencies, which are greater than 120 Hz for fruit flies and blowflies, and far away from the frequency band of the head movements we observed (Rosner et al., 2009; Van Hateren et al., 1999). Spontaneous head saccade frequency also moderately decreased when flies were presented with a uniform background, consistent with previous work (Wolf and Heisenberg, 1980) suggesting that overall background luminance and/or visual features influence the trigger of spontaneous head saccades (Figure Supplement 5E). Taken together, we show that even in the absence of visual motion, fly eyes are not still and are punctuated by rapid spontaneous movements. These movements are consistent with illusory motion in the absence of external motion (Salem et al., 2020). Although we cannot infer the functional role of these head movements with confidence, they may be part of a search strategy for active vision systems and may even serve to refresh visual inputs in a way similar to human micro-saccades (Martinez-Conde and Macknik, 2017).

### Head movements are gated by behavioral state

For faster visual stimuli, the head reached its anatomical limits more quickly during smooth intervals (Figure 4A,B). Therefore, reset saccade rate would presumably increase with stimulus speed, however intriguingly this was not always the case at higher speeds (Figure 2B). Flies would not always perform a head saccade immediately when the anatomical limit was reached (Figure 4A,D). Thus, flies did not reduce retinal slip during these periods as the head remained near the edge of the thorax. But flies are physically capable of robust gaze stabilization enabled by reset saccades, even at the highest stimulus speed presented in this study (Video 4). So, what could be causing flies to keep their heads at their anatomical limit instead of performing head saccades to reset gaze direction?

We noted that flies would occasionally extend their legs, indicative of a fictive landing response (Ache et al., 2019; Borst, 1986; Goodman, 1960; Tammero and Dickinson, 2002b; van Breugel and Dickinson, 2012) (Figure 5A, Video 7). Leg extension occurred more frequently at higher stimulus speeds, consistent with work showing that flies switch from forward flight to expansion avoidance based on the rate of expansion optic flow (Figure 5B) (Liu et al., 2019; Reiser and Dickinson, 2013). Here, however, we show that flies can switch to a landing state based on the rate of *rotational* optic flow. Interestingly, during leg extension, flies seldom generated head reset saccades, and the head consequently remained near its anatomical limit (Figure 5C). We hypothesized that when flies enter a landing state, head movements may be modulated differently. To test this hypothesis, we used machine vision to detect the leg extension reflex (see Methods). Using the tracked leg kinematics, we classified bouts of flight into binary states: stabilizing flight and landing, and pooled head movements across these states. Less than 5% of all head saccades occurred when flies were extending their legs (*N* = 3,571 saccades, same as 30° wavelength data from Figure 2) (Figure 5C). Correspondingly, during leg extension the head operated at overall lower angular velocities and greater excursion, consistent with the head remaining at its anatomical limit instead of performing a reset saccade (Figure 5D, Video 8). These results provide a possible explanation for the reduced influence of stimulus speed on head saccade rate at higher speeds (Figure 2B): flies were operating in a different behavioral state under these conditions. Furthermore, these results imply that flies can escape the inner-loop head gaze stabilization reflex active during forward flight, via an outer-loop initiation of landing and suppression of head reset saccades (Hardcastle and Krapp, 2016). Altogether, the behavioral state of the fly gates stabilizing head movements and saccades.

**Figure 5.**
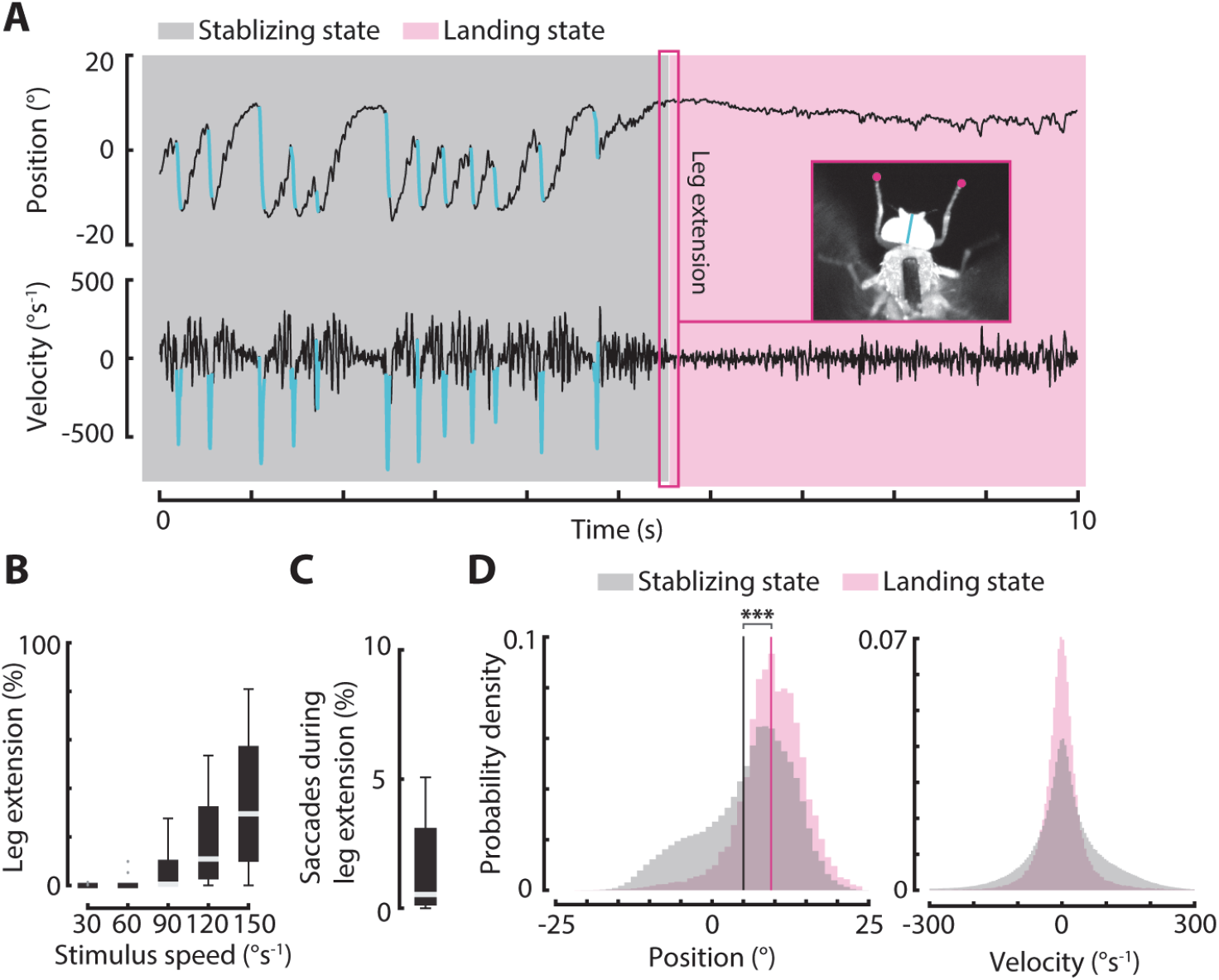
Head movements are gated by behavioral state. **A)** Example trial for −120°s^−1^ stimulus motion at 30° spatial wavelength (Video 3). Pink background: flies performing gaze stabilization via a combination of smooth and saccadic head movements. Grey background: flies initiating a landing or collision avoidance response, as classified by leg extension (inset). Cyan traces: head saccades. **B)** Percent of time flies spent extending their legs vs. stimulus speed. Flies rarely extended their legs at lower speeds, so the upper and lower boxplot quartiles are equal to the median (0%). Distributions represent the means for each fly at each stimulus speed. Grey markers: outliers. **C)** Distribution of saccades occurring during leg extension divided by the total number of saccades across all stimulus speeds. **D)** Distributions of head angular positions and velocities while stabilizing (no leg movements) and landing (leg extension) across all stimulus speeds. Counter-clockwise intervals were inverted and pooled with clockwise intervals. *t*-test, *** = *p* < 0.001. 30° spatial wavelength: *n* = 12, *N* = 4,091 saccades.

### The head saccade trigger implicates proprioceptive feedback

When flies are actively engaged in gaze stabilization, i.e. not extending their legs, most evidence points to the angular position of the head playing the largest role in triggering a saccade (Figure 2C, 4A). Therefore, proprioceptive feedback could drive the initiation of head saccades. Specifically, flies could sense the angular position of their head, and when the head nears the anatomical limit of the head-thorax joint, a neural command of mechanosensory origin could be sent to trigger a reset saccade. The cumulative distribution function of saccade trigger positions suggests that the mechanism consists of a soft threshold ~9° from the head neutral position (Figure 6A). Based on these findings, we hypothesized that saccade rate would be influenced by the operating range of the head; specifically, flies would perform saccades less frequently for visual stimuli that did not drive the head near its anatomical limits. To test this hypothesis, we provided flies with wide-field, sinusoidal visual motion at combinations of five amplitudes spanning 3.75°– 18.75° and six frequencies spanning 0.5–12 Hz corresponding to distinct mean speeds (see Methods) (Figure 6B, Video 9–12). The head response was tuned to each distinct combination of stimulus amplitude and frequency, leading to distributions of head positions of varying widths (Figure Supplement 6A). The distributions were correlated with the gain of head movements during smooth movements (Cellini and Mongeau, 2020b). Generally, stimuli with larger amplitude and lower frequency elicited head movements with the widest distributions, whereas lower amplitude and higher frequency combinations generated the narrowest distributions (Figure Supplement 6A). We parameterized each distribution by twice the mean of its absolute value, which approximated the operating range of the head (Figure 6B,C, Figure Supplement 6A).

**Figure 6.**
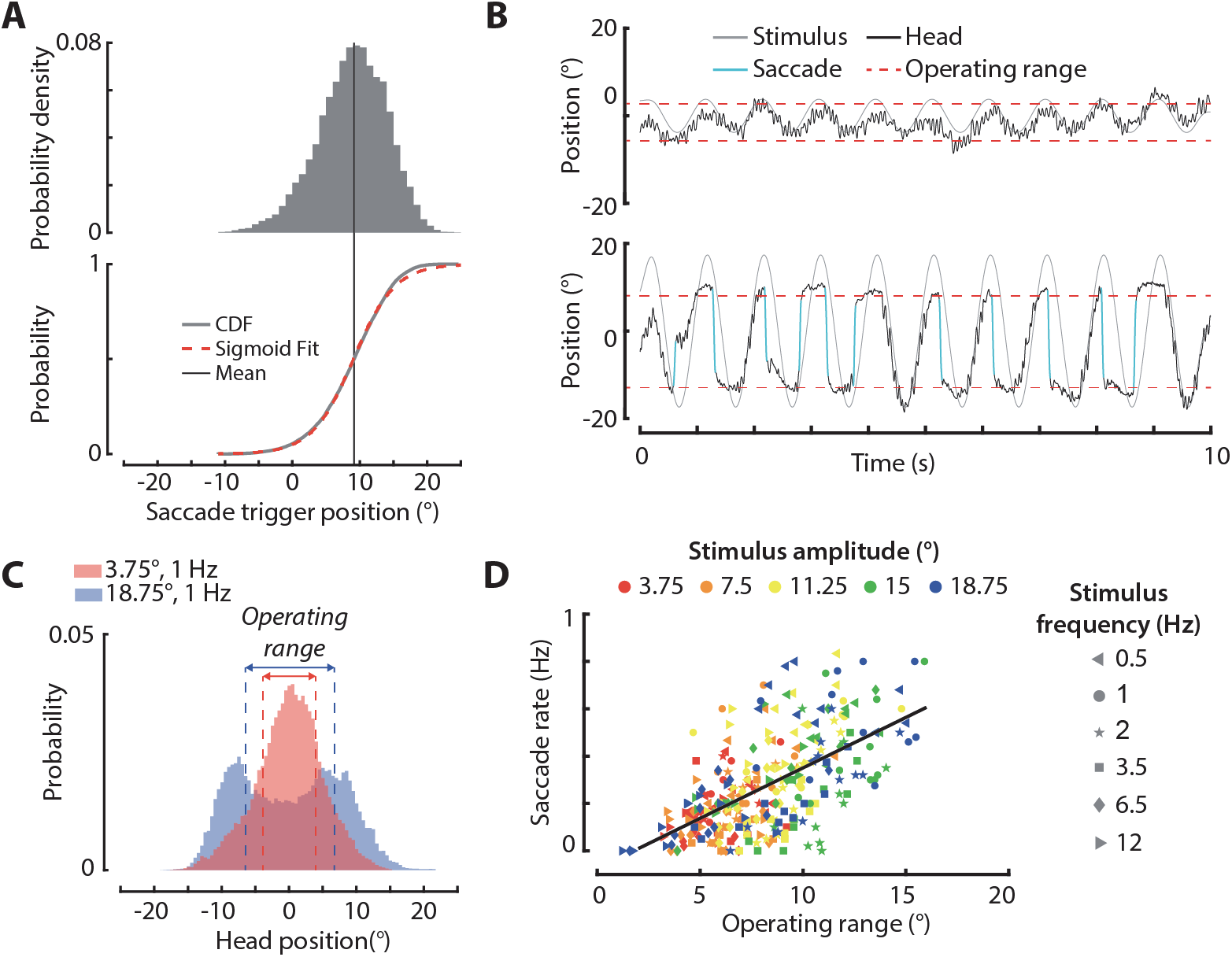
The head saccade trigger implicates proprioceptive feedback. **A)** Distribution of head saccade trigger positions (top, same as Figure 2C) and the corresponding cumulative distribution function with a sigmoid fit (bottom; see Methods). **B)** Example trials for sinusoidal stimulus motion at 1 Hz oscillation frequency with an amplitude of 3.75° (top) and amplitude of 18.75° (bottom). Red dashed lines: range of head movements for each stimulus. **C)** Distributions of head angular positions for 1 Hz with 3.75° and 18.75° amplitude sinewave visual stimuli. **D)** Correlation between head operating range and saccade rate (r = 0.61, *p* < 0.001). Each data point represents an individual fly’s response at a given stimulus amplitude-frequency combination. 3.75°: *n* = 8 flies, *N* = 746 saccades. 7.5°: *n* = 10 flies, *N* = 602 saccades. 11.25°: *n* = 11 flies, *N*= 957 saccades. 15°: *n* = 10 flies, *N* = 1,068 saccades. 18.25°: *n* = 11 flies, *N* = 1,057 saccades.

We discovered that the operating range of the head had a strong influence on saccade rate (Figure 6D, Figure Supplement 6B). While the oscillatory stimuli elicited less saccades overall than the ramp stimuli, we observed a strong correlation between the head’s operating range and saccade rate (r = 0.61, *p* < 0.001). Neither the stimulus amplitude (r = 0.29), frequency (r = 0.34), or mean speed (r = 0.21) were correlated as strongly with saccade rate as the operating range of the head. These results demonstrate the role of head orientation, as opposed to visual information, in triggering head saccades. Effectively, a larger operating range increased a fly’s chance of moving its head to its anatomical limit, and thus increased the probability that a saccade was triggered. The stimuli corresponding to the narrowest distributions of head movements still elicited some saccades, but at a rate similar to spontaneous saccades rates (Figure Supplement 6B). This point was further supported by comparing the head orientation at saccade initiation across stimuli. The majority of saccades initiated within an overall narrow operating range were triggered nearer to the head center line, similar to the start orientation for saccades performed for a static background (Figure Supplement 6B; Figure 2C). On the other hand, saccades for wider head angular distributions were triggered near the head’s anatomical limit (Figure Supplement 6B). These head saccades usually preceded changes in stimulus velocity, confirming that they were indeed “reset” saccades (Figure 6B, Video 10). Taken together, these results support the hypothesis that the activation of head saccades is based on a position-based threshold. This mechanical threshold is stochastic and is likely influenced by spatiotemporal visual information as well as endogenous stochastic processes. However, our results provide strong evidence in favor of proprioceptive feedback as the driving mechanism in head saccade activation.

### The head and wings switch between smooth and saccadic control simultaneously

If the origin of head reset saccades is truly based on proprioceptive information from the neck sensory system, how does this trigger influence body saccades (via wing saccades) and are head and body saccades always coupled? Indeed, a head-based proprioceptive trigger might suggest that the head could perform reset saccades independently of the body. However, prior work showed that head and body saccades are coupled in free flight (Schilstra and van Hateren, 1998) and in magnetically tethered paradigms (Kim et al., 2017), although these studies analyzed spontaneous saccades exclusively. Smooth head and wing movements in rigidly tethered flight are also coupled when presented with a moving panorama (Cellini and Mongeau, 2020b; Duistermars et al., 2012; Kim et al., 2015), but can be decoupled about yaw for simultaneous small- and wide-field motion (Fox and Frye, 2014), thereby suggesting that a parallel visual pathway may elicit wing steering responses. To elucidate head-wing interactions, we tracked wingbeat kinematics and compared them to head kinematics. In the rigid tether paradigm, the difference between left and right wingbeat amplitude (ΔWBA) is roughly proportional to the yaw torque acting on the body (Tammero et al., 2004) (Figure 1A). Therefore, saccadic wing movements, i.e. hitches in ΔWBA, indicate fictive body saccades.

We developed an algorithm to detect and extract wing saccades from ΔWBA traces (see Methods) and then classified head saccades as either 1) having an accompanying wing saccade within a 100 ms window or 2) not having an accompanying wing saccade (Figure 7A, Video 13). About 50% of head reset saccades had an accompanying wing saccade, but some head reset saccades occurred without a clearly identifiable wing saccade (Figure 7A, Figure Supplement 7A). However, wing movements around head saccades without an accompanying wing saccade still showed a general positive correlation with head movements (Figure Supplement 7B). The lesser peak velocity and amplitude of these wing movements rendered detection highly challenging, as they were not sufficiently above the noise floor. This was especially problematic in trials with a high saccade rate (Video 4). Wing saccade detection parameters could be tuned to detect these wing hitches, with the caveat of an increased false-positive rate (see Methods). Although we could not achieve a one-to-one detection between head and wing saccades, the general trend of ΔWBA around all head saccades indicates that head and wings perform reset saccades concurrently with the wing response leading the head by ~5–10 ms (Figure 7B,C). Flies did not actively ‘brake’ via a negative ΔWBA signal to terminate saccades and instead followed a trajectory closely mirroring the head.

**Figure 7.**
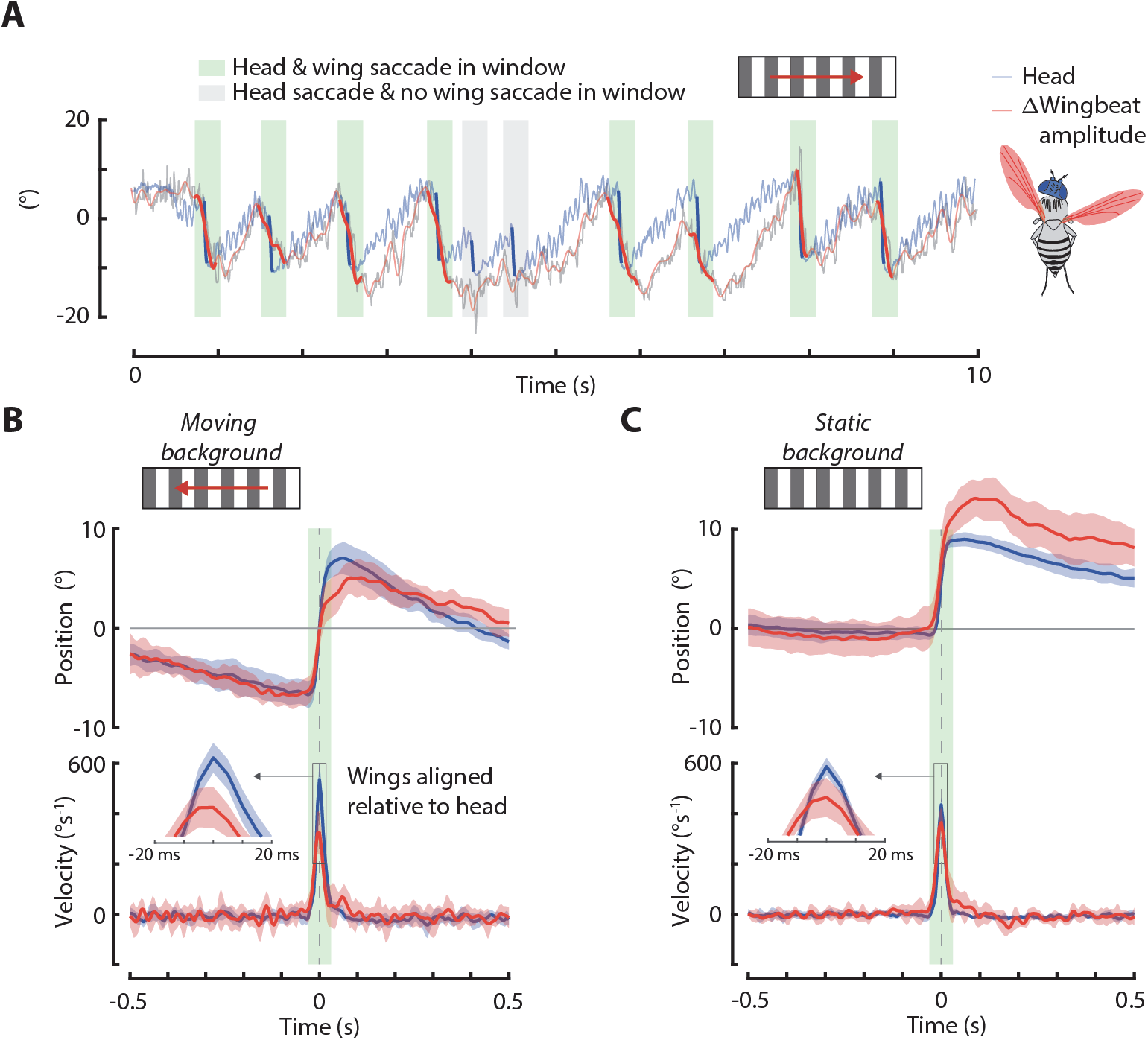
The head and wings switch between smooth and saccadic control simultaneously. **A)** Example trial showing the head (blue) and ΔWBA (red) response to a stimulus moving at 30°s^−1^ with a 30° spatial wavelength. Saccades are highlighted. Green area: a head and wing saccade were detected within a 100 ms interval. Grey area: only a head saccade was detected. **B)** Head position and velocity aligned to head saccade velocity peak time and ΔWBA position and velocity aligned relative to the head, such that the true time difference between head and ΔWBA was maintained. Data for synchronous head-wing saccades (green window) is shown. Inset: relative difference in the peak times of head and wing saccades. Each ΔWBA trace was normalized to the mean of the first and last 300 ms. Counterclockwise saccades were inverted and pooled with clockwise saccades Shaded regions: ±1 standard deviation. **C)** Same as B) but for saccades detected during the presentation of a static background. Each ΔWBA trace was normalized to the mean of the first 300 ms. 30°s^−1^ background: *n* = 12 flies, *N* = 829 head saccades. Static background: *n* = 8 flies, *N* = 359 saccades.

Interestingly, head and wing saccades lasted for a similar duration of ~50 ms, as evidenced by the clear switch from saccadic to smooth control around each head saccade (Figure 6B,C, Figure 4A). This is in contrast to prior work suggesting that wing saccades last upwards of 500 ms in rigidly tethered flies (Heisenberg and Wolf, 1986; Wolf and Heisenberg, 1980). This discrepancy spawns from conflicting definitions of the time scale of wing saccades. Prior studies have classified spontaneous wing saccades as long duration spikes in ΔWBA, consisting of an initial spike in ΔWBA followed by a slow decay back baseline. We observed these same ΔWBA spikes in flies presented with a static background (Figure 7C, Figure Supplement 5A, Video 5). However, using raw ΔWBA to classify fictive body saccades in rigid tether paradigms is problematic, as the lack of reafferent feedback—presumably from the halteres—could artificially exaggerate the transient response (Bender and Dickinson, 2006; Dickinson and Muijres, 2016). We contend that wing saccades should be classified by the rate of change of ΔWBA (see Methods), therefore only the initial spike above baseline ΔWBA should define saccade duration. This is a more precise method of classifying flight into smooth and saccadic bouts, in addition to being more consistent with the duration of body saccades in free flight (Muijres et al., 2015). If wing saccades did truly last 500 ms, then this would imply that the head and wings were often operating under different control (smooth *vs*. saccadic), which did not seem to be the case here. Taken together, head and wing reset saccades are under simultaneous control and have similar durations in the rigid tether.

### Proprioceptive feedback influences both head and wing saccade initiation

Thus far our results suggest that 1) head saccades are triggered primarily by proprioceptive feedback and 2) head and wing saccades are triggered concurrently. But does proprioceptive head feedback directly influence wing saccade control? For a purely vision-based saccade trigger, one might predict that head-fixed flies perform saccades *more* frequently, due to the lack of stabilizing smooth head movements resulting in increased retinal slip error. However, based on our findings, we hypothesized that wing saccade rate in head-fixed flies would be less than that in head-free flies, due to the head never reaching its anatomical limit. To test this hypothesis, we presented head-fixed flies with the same ramp stimuli presented to head-free flies and compared the overall ΔWBA response and wing saccade rate (Figure 8, Video 13). Head-fixed flies exhibited a greater mean wing response with less intra- and inter-trial variance than head-free flies (*t*-test, *p* < 0.001)(Figure 8A,B). Without reset saccades, we would expect the wing response to be larger and less variable, thus these results suggest that wing saccades in head-fixed flies occur less frequently, if at all, and/or with reduced amplitude. To confirm this, we culled wing saccades from head-free and head-fixed ΔWBA traces and found that head-fixed flies did indeed perform reset saccades less frequently (*t*-test, *p* < 0.001) (Figure 8C). Wing proprioceptive feedback could also influence the reset saccade trigger. In the rigid tether, restricting body motion causes the ΔWBA signals to operate in a broader range than in free flight (Dickinson and Muijres, 2016)(see Video 4).

**Figure 8.**
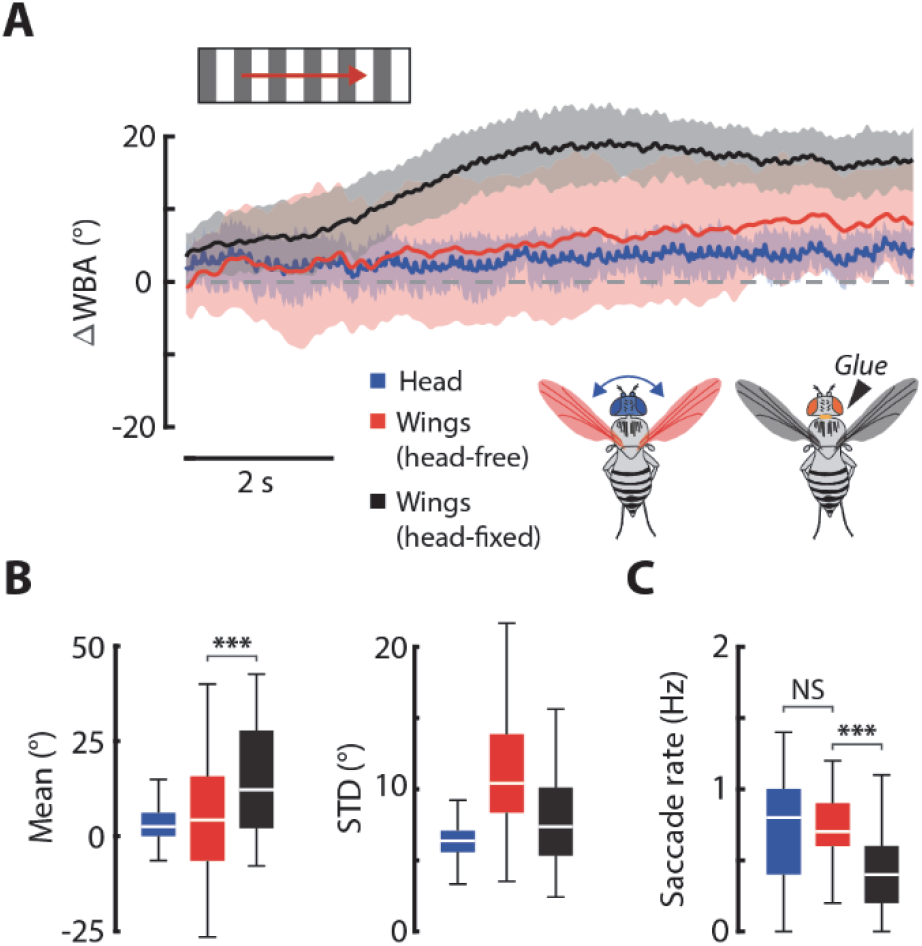
Proprioceptive feedback influences both head and body saccade initiation. **A)** The mean response of the head (blue) and ΔWBA (red) for head-free flies, along with the mean ΔWBA (black) for head fixed flies, presented with a 30°s^−1^ stimulus at 30° spatial wavelength. Shaded regions: ±1 standard deviation. **B)** The distributions of intra-trial means and standard deviations for head-free and head-fixed flies. Colors follow same scheme as in A). **C)** Saccade rates for the head and wings in head-free flies for a 30°s^−1^ stimulus as well as wing saccade rates for head-fixed flies for the same stimulus. Same color scheme as in previous panels. *t*-test, *** = *p* < 0.001. Head-free: *n* = 12 flies, Head-fixed: *n* = 12 flies

We hypothesized that a threshold in wing angular excursion could provide a signal to initiate a reset saccade. To uncover the possible role of wing proprioceptive feedback, we compared the ΔWBA distributions at the start and end of a saccade in head-free flies (Figure Supplement 8). In addition, we compared saccade start and end distributions for the left and right WBA separately (Figure Supplement 8). Compared to head saccade start and end distributions (Figure 2C), the wing start and end distributions significantly overlapped in all cases and were not distinct from the ΔWBA distribution, suggesting that neither absolute WBA nor ΔWBA may provide a robust trigger signal for saccade initiation. Indeed, absolute wingbeat amplitude can vary substantially in flies and therefore may not provide a salient trigger signal (Duistermars et al., 2007). In conjunction with our prior findings, these results strongly implicate proprioceptive feedback of head orientation as the primary origin for the trigger of both head *and* wing reset saccades.

## Discussion

By combining smooth head movement and reset saccades, flies stabilize gaze in a punctuated manner, overcoming anatomical limits in a way analogous to vertebrate eye reflexes. Head reset saccades were strongly stereotyped and interrupted smooth movement gaze stabilization for as little as 50 ms, representing less than 5% of total flight time (Figure 2). Flies leveraged the elasticity of the neck joint to initiate head reset saccades, demonstrating the role of passive mechanics in head control (Figure 3). Head saccades enables flies to quasi-continuously stabilize visual motion, reducing retinal slip by up to 70% between head saccades (Figure 4). Reset saccades were initiated when the head was near its anatomical limits and their rate was highly dependent on the operating angular range of the head, implicating proprioceptive feedback in saccade initiation (Figure 2C, 6). Visually and spontaneous head saccades were triggered at different absolute head positions, suggesting different trigger origins (Figure 2C). Higher stimulus speeds elicited a landing response, which suppressed head smooth movements and saccades, demonstrating that behavioral state gates head visuomotor responses (Figure 5). Lastly, the head and wings switched between smooth and saccadic control concurrently but fixing the head significantly reduced wing saccade rate (Figure 7,8). These results further support the role of proprioceptive feedback in triggering reset saccades. To our knowledge, we demonstrate for the first time that wing control can be directly influenced by head movements. Taken together, our work demonstrates the critical function of a hybrid strategy for visuomotor control that provides a robust solution to overcome sensor saturation limits.

### Reset saccades as a hybrid control solution for systems with saturation nonlinearities

In the task studied here, the dynamics of the head are inherently nonlinear due to the anatomical range of the joint. In control theory, this class of nonlinearity is categorized as a *saturation nonlinearity*. Essentially, when the head is saturated (oriented at anatomical limit; Figure 4A), no matter how much control effort (torque at neck joint) is applied, the head will remain in this state. Interestingly, if a fly’s stabilization control strategy were purely linear, the head would saturate quickly and remain at its anatomical limit, rendering further efforts to stabilize gaze futile (Figure 4A,B). We confirm that flies solve this problem by performing a rapid head and wing reset saccade when the head nears this saturated region, allowing stabilizing movements to begin anew (Figure 2C, Video 1). Because reset saccades lasted ~50 ms, flies experienced only a momentary disruption of smooth gaze stabilizing movements. Overall, flies were able to spend ~95% of the time performing stabilizing head movements. From a design perspective, it would be optimal for flies to reset their head orientation as fast as possible so as to minimize the periods where gaze is not stabilized, which is consistent with the constant rapid saccade dynamics we observed across stimulus speeds. The switch from smooth to saccadic control comprises a hybrid control scheme that allows flies to deal with the head’s saturation nonlinearity while still performing the given task (gaze stabilization) (Figure 9A). This concept can be illustrated by artificially removing saccades from a fly’s head and wing trajectories (see Methods) and observing the continuous stabilization response (Figure 9B). Altogether, the hybrid mechanism allows the smooth controller to operate in the region where it is most effective (i.e. not saturated). Crucially, this strategy only applies to a velocity sensitive control system like the gaze stabilization reflex, as opposed to a position sensitive one like object localization, where reset saccades would result in swiftly increasing error. Flies seldom perform reset saccades during object tracking tasks, thus supporting this notion (Mongeau and Frye, 2017).

**Figure 9.**
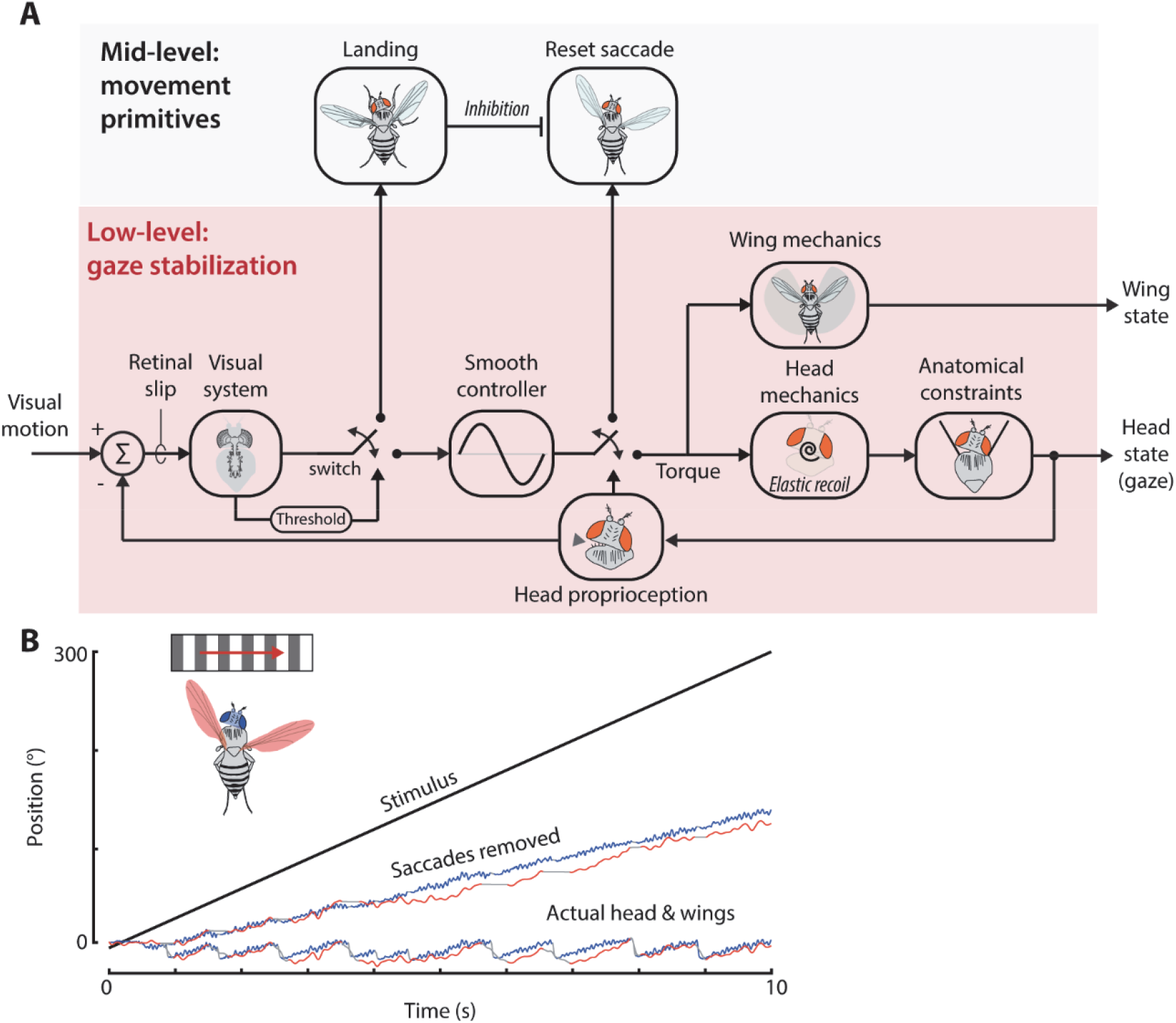
Summary of findings. **A)** Proposed control scheme describing how reset saccades are triggered. The velocity-based stabilization reflex (low-level) is switched-off by an outer-loop saccade motor command when a fly senses that its head is oriented near its anatomical limit. The fly presumably senses its head orientation via a proprioceptive mechanism. The saccade motor commands result in a rapid reset of head and wing position, leveraging the passive elasticity of the neck joint. Head orientation feeds back to alter the fly’s perceived visual motion (retinal slip). The fly continuously monitors retinal slip and is more likely to exit the stabilization reflex via an outer-loop initiation of landing for higher retinal slip speeds. Landing inhibits the saccade motor program. **B)** Using an example flight trial for continuous visual motion, here we show the head (blue) and wing (red) responses with saccades artificially removed and the smooth intervals interpolated together (see Methods). The actual head and wing responses are shown for comparison. Saccades and the corresponding interpolated data are highlighted in grey. The stimulus is shown for comparison (black).

### Proprioceptive mechanisms

The consistent head orientation at the time of saccade initiation, the influence of the head’s operating range on saccade rate, and the decrease in saccade rate in head-fixed flies strongly suggest that reset saccades are triggered by proprioceptive feedback (Figure 2C, 6, 8). Flies most likely monitor the orientation of their head and initiate a reset saccade when the head reaches some predefined threshold near the anatomical limit (Figure 8). Neural recording within the neck motor system of soldier flies showed that a pair of mechanosensory hair fields located on the prothorax, the prosternal organs (PO), encode information regarding head posture about yaw, pitch, and roll (Paulk and Gilbert, 2006). However, in blow flies, unilaterally shaving the PO results in a postural bias primarily about the roll and pitch axes, with less of an influence observed in yaw (Preuss and Hengstenberg, 1992). This result suggests that there may be other mechanosensory mechanisms involved in the control of head yaw. Flies have additional mechanoreceptors that may be involved in head control, including the prothoracic chordotonal organs (CO) and several larger hairs on the back the head (Strausfeld et al., 1987). Altogether, reset saccades are most likely triggered by the stimulation of mechanosensory organs at the base of the fly neck. Our work provides testable hypotheses concerning the physiological mechanisms involved in triggering reset saccades.

### Potential for a conserved mechanism across species

In addition to fruit flies, blow flies (Land, 1973), hover flies (Collett and Land, 1975), locusts (Kien and Land, 1978), beetles (Varju and Bolz, 1980), and mantids (Lea and Mueller, 1977; Liske and Mohren, 1984) perform reset saccades of the head and/or body during visuomotor tasks. Presumably, other insects have a similar proprioceptive mechanism that enables this behavior. The locusts’ neck sensory system contains the myochordotonal organs (Shepheard, 1973), similar to the CO in flies, which have a strong influence on head saccade during yaw gaze stabilization. Interestingly, reset saccades virtually disappear when these organs are cut, suggesting that a proprioceptive mechanism may be necessary (Shepheard, 1974). A similar concept applies to mantids, where the reset saccade rate in response to a moving target is nearly cut in half when hairs at the base of the neck are shaved, and saccade speed and amplitude increase (Liske and Mohren, 1984). Intriguingly, the idea of proprioceptive triggered reset saccades may span multiple taxonomic classes, as eye movements in crabs (Horridge, 1967) and primates (Cheng and Outerbridge, 1974) display remarkably similar movement patterns. Indeed, mammal eyes have multiple proprioceptive mechanisms that could influence saccade control during optokinetic nystagmus (Donaldson, 2000), however, to our knowledge the mechanisms underlying the trigger of reset saccades during optokinetic nystagmus is not known in humans and non-human primates. Altogether, a proprioceptive saccade trigger may be a conserved characteristic of active vision systems.

### Head-wing interactions

To our knowledge, we demonstrated for the first time that head orientation influences wing saccades (Figure 8), establishing a causal link. Prior work has shown that 93% of saccades in free flight can be linked to salient visual features in a fly’s environment (Censi et al., 2013). The remaining saccades were attributed to unobservable internal states or spontaneous behavior. Intriguingly, the reset saccades studied here could provide a possible explanation for a portion of these “spontaneous” saccades. We propose that some of these saccades could be triggered by head saturation, presumably measured by the PO or CO, or even by stochastic processes underling proprioceptive feedback from hair plates. Smooth wing steering gain and coordination is also influenced by head movements during gaze stabilization tasks (Cellini and Mongeau, 2020b). Furthermore, fixing the head of a fly at an angle in complete darkness results in a wing steering response in the same direction of the head, demonstrating that head orientation alone (without visual influences) affects wing control (Liske, 1977). These studies, in conjunction with our findings, present an avenue for future work that investigates links between proprioceptive feedback from the neck sensory system and flight control.

### Neural control of saccades

Our results pose testable hypotheses concerning how reset saccades might be initiated in the fly nervous system. The first possibility is that neck proprioceptors (PO, CO, etc.) sense when the head is near its anatomical limit which triggers coordinated head and wing reset saccades via direct, descending pathways to the head and wing motor centers. Head and wing neuropils are innervated by many mutual descending neurons (DN) and head position influences wing steering efforts in flies, thus the head-neck sensory system could directly influence *both* head and wing control (Cellini and Mongeau, 2020b; Liske, 1977; Namiki et al., 2018). At present, it is not known whether the saccade motor program is initiated peripherally or from more central brain processes. Our results suggest that head angular position is linked to the trigger of reset saccades, but because wing saccades precede head saccades, the saccade motor program may be located more centrally (Figure 9A). We believe this is the more likely scenario because wing saccades precede head saccades by ~5 ms (Figure 7). If saccades were triggered via direct pathways from the neck that bypass the central brain, it would be reasonable to expect that head saccades would precede wings saccades due to the shorter distance signals would have to travel. Furthermore, motor-related efference copies act on peripheral visual interneurons during fictive saccades, suggesting that these saccade-related signals originate from more central brain processes (Kim et al., 2017, 2015). Taken together, our work suggests that saccades may originate from a motor program located more centrally within the nervous system of *Drosophila*.

### Effect of behavioral state on head movements

We found that coordinated smooth and saccadic head and wing movements are an integral component of the gaze stabilization system but that these sensorimotor processes are not always active (Figure 5). For high retinal slip speeds flies shifted to a landing state which significantly reduced head smooth movements and saccades (Figure 5A). Flies rarely performed reset saccades during landing and instead maintained a persistent head orientation biased towards one side, in the direction of the visual stimulus (Figure 5C,D). This implies that the behavioral state is somehow inhibiting the saccade command center. At present, it is unclear how landing might modulate visuo-motor processing. A possibility is that the saccade trigger is influenced by neuromodulators acting at the periphery that inhibit neck sensory afferents. The response range, gain, and latency of visual processing in flies is also influenced by state-dependent neuromodulation, which could play a role in the suppression of head movements (Ache et al., 2019; Longden and Krapp, 2009; Maimon et al., 2010). Why head movements are reduced during landing is intriguing, as corrective head movements would presumably be critical during this task (Liu et al., 2019). However, restricting body movements may complicate the functional interpretation of head saccade suspension during landing. One possibility is that flies were attempting to perform a co-directional turn in the direction of visual motion which, because of the rigid tether, simply resulted in a continuously saturated head position during landing.

### Effects of restricting body motion

Our results show that rigidly tethered flies regularly perform reset saccades (Figure 2), but it remains unclear whether head reset saccades are common in free flight where the body is free to move. It could be that reset saccades are less prominent in freely flying flies due to body movements that reduce the probability of the head reaching a saturated state. Furthermore, body movements can rapidly reduce retinal slip, thereby in free flight head movements might be very small to compensate for similarly small visual drift (Cellini and Mongeau, 2020b; Van Hateren et al., 1999). However, over time, the accumulated drift may be sufficient to generate a reset saccade via spatiotemporal integration of visual and/or mechanical origin, as demonstrated in a magnetic tether (Mongeau and Frye, 2017). Unlike the head, the body has no anatomical constraints restricting its range of motion, thus flies could stabilize gaze via a combination of head and body movements that would seldom require a reset saccade. Nevertheless, reset body saccades have been observed in freely flying hover flies, demonstrating that this behavior is not limited to rigidly tethered flight (Collett and Land, 1975). Furthermore, reset body and head saccades are performed by magnetically tethered flies, where the body is constrained to yaw but the head is free to move (Land, 1973; Mongeau and Frye, 2017). In response to wide-field visual motion, magnetically tethered flies performed reset saccades with a rate of ~0.15 Hz, more than five times less than the rate reported here, although this result was dependent on stimulus speed (Mongeau and Frye, 2017). This suggests that reset saccades are activated less often in flies with unrestricted body dynamics. Nevertheless, the consistent head orientation of magnetically tethered blowflies at the instance of reset saccades (~10–15° from midline) supports our proposed proprioceptive-based trigger (Land, 1973)(Figure 9A). Moreover, prior studies showed that co-directional body saccades in magnetically tethered and freely flying fruit flies are immediately followed by a rapid anti-directional head movement. Although these studies did not interpret this anti-directional movement as a separate saccade, the dynamics suggest that it is indeed a reset saccade, presumably executed to recenter the head to body midline (Kim et al., 2017; Van Hateren et al., 1999). An exciting avenue for future research will be to elucidate the role of the head and its influence on flight control in paradigms with unrestricted body dynamics.

### Bio-inspired applications

With a growing interest in bio-inspired technology, many recent studies have focused on the development of vision sensors inspired by insect eyes for autonomous vehicles (Serres and Viollet, 2018). A particular focus has been placed on understanding and instantiating the physiological and motion processing mechanisms that make insects such proficient flyers. However, less emphasis has been placed on one of the most defining features of insect vision: insect eyes move via head movements, and are thus active visual sensors (Bajcsy, 1988). As current micro-air vehicles have recently begun to incorporate movable sensors, roboticists and engineers should consider saccade-inspired hybrid control for vision systems with saturation nonlinearities (Iyer et al., 2020). For example, a recent study developed a wireless steerable vision sensor for insect-scale flying systems (Iyer et al., 2020). This sensor can be actuated through ~60° in yaw via a piezo-based actuator, and thus has saturation constraints analogous to the mechanics of an insect head. In order to continuously stabilize gaze, this sensor’s movement could be programmed to perform reset “saccades”. A proprioceptive sensor, analogous to the PO or CO in flies, would be all that is required to implement this control scheme. Altogether, this hybrid control strategy has important applications for active vision systems in insect-scale robots and beyond that must overcome limits in sensor range of motion.

## MATERIALS AND METHODS

### Animal preparation

We prepared the animals for each experiment according to a protocol that has been previously described (Reiser and Dickinson, 2008). Briefly, we cold-anesthetized 3–5 day old females flies (wild-type *Drosophila melanogaster*) by cooling them on a Peltier stage maintained at approximately 4°C. We fixed flies to tungsten pins (A-M Systems) by applying a small drop of UV-activated glue (XUVG-1, Newall) to the thorax. The pin was placed on the thorax projecting forward at an angle of approximately 90°. For head-fixed experiments, a small drop of glue was placed atop the neck joint. After a minimum 30-minute period of recovery, flies were used for experiments.

### Experimental protocol

The rigid-tether arena has been described elsewhere (Reiser and Dickinson, 2008). Briefly, the display consists of a cylindrical array of 96 × 32 light emitting diodes (LEDs, each subtending 3.75° on the eye) that wrap around the fly, subtending 330° horizontally and 112° vertically (Figure 1A). Flies were rigidly mounted to a rod inside the arena and illuminated from above with a single 940 nm LED (Figure 1A). Flies were attached to a rigid rod and placed in the arena at an angle near the natural flight angle (~45° in pitch). The position of the fly was adjusted until the fly was in the frame, and in focus of the camera. Flies then underwent a closed-loop bar fixation period of approximately ten seconds to ensure that the wings were being accurately measured and that the flies were not visually impaired. Flies that failed to fixate the bar were not used for experiments. All experimental trials were interspersed with closed-loop bar fixation bouts of five seconds to ensure attentiveness throughout the length of the experiment. We recorded videos of the fly from above at 200 frames per second (fps) with an infrared-sensitive camera (acA640-120gm, Basler). During trials, video frames were collected using an external trigger, and synced to the start of visual motion. We acquired signals (stimulus position and camera trigger) at 5 kHz.

### Visual stimuli

In order to study the dynamics of the hybrid visual system of flies, we designed a visual stimulus that elicited both smooth and saccadic head movements in yaw. Previous work showed that oscillatory wide-field movements with smaller amplitudes and intermediate velocities elicit mostly smooth head movements in rigidly tethered flies (Cellini and Mongeau, 2020b). Although there is little prior work investigating head saccade activation in rigidly tethered flies, body saccades have been shown to occur frequently for constant velocity wide- and small-field motion (Mongeau and Frye, 2017). Therefore, we used constant velocity, wide-field visual stimuli, which proved sufficient to elicit robust smooth and saccadic movements. Individual flies were presented with velocities of 30, 60, 90, 120, 150 °s^−1^ in both directions at fixed spatial wavelengths and in randomized order. The spatial wavelengths of the panorama presented were 22.5°, 30°, 60°. All stimuli were presented at the maximum contrast offered by the LED arena, near a 100% Michelson contrast. When investigating spontaneous saccades, we displayed a stationary panorama at the same wavelengths and contrasts as the moving stimuli but added a all-on, all-off, and randomized spatial wavelength grating.

To investigate how the operating angular range of the head affected head saccades, we employed sinewave visual motion at amplitudes of 3.75°, 7.5°, 11.25°, 15°, and 18.75° at frequencies of 0.5 Hz, 1 Hz, 2 Hz, 3.5 Hz, 6.5 Hz, and 12 Hz. These visual stimuli have been further described in past work exploring smooth head movements in *Drosophila* (Cellini and Mongeau, 2020b). Sinewave motion can be described using the mean speed, *s_mean_* = 4*Af*, where *A* is the amplitude of the sinewave and *f* is the frequency of oscillation. We use this metric to compare stimuli with similar mean speeds, but with different amplitudes and frequencies.

### Kinematic measurements

We used custom computer-vison software (MATLAB, MathWorks, Natick, MA, USA) to track the angle of the head relative to the body and left and right wingbeat amplitude (WBA) in units of degrees. Briefly, we manually defined the head rotation point on the neck and the initial angle of the head. Then we automatically tracked features on the fly head (base of the antennae) and measured the change in angle of the feature relative to the rotation point for every frame. We also measured WBA in each frame using an edge detection algorithm similar to Kinefly (Suver et al., 2016). Because flies flapping frequency exceeds 200 Hz, the leading edge of the blur of each wing approximates WBA.

Head roll was measured using the relative width of the left and right eye in every video frame (Kim et al., 2017) (Video 2). The lighting for these experiments was adjusted to ensure a clear distinction between the fly head and compound eyes. In short, we stabilized recorded frames in the head yaw reference frame using our yaw measurements. Then we measured the pixel intensity in a region surrounding the center of the head along the horizontal axis. The mean intensity displayed two distinct peaks representing the left and right eyes, where the width of the peaks corresponded to the width of each eye in pixels. We computed the roll index, defined as

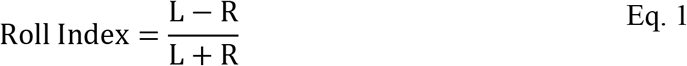

where *L* and *R* are the widths of the left and right eye, respectively. This index is linearly related to the true head roll by a constant calibration factor in tethered flies (Kim et al., 2017). We used this calibration factor to approximate head roll in a similar experimental paradigm.

To tack the legs, we trained an artificial neural network using the pose estimation Python library DeepLabCut (Video 7) (Nath et al., 2019). We used a training dataset of ~1,500 frames of labeled images consisting of the left and right front legs, and a test dataset consisting of ~500 labeled frames. After training for more than 100,000 iterations we achieved a test accuracy within 3 pixels. Because our analysis was not concerned with the precise movements of the legs, we classified frames as either having 1) leg extension or 2) no leg extension. We then applied a median filter to the binary classification for all the frames in each experiment to remove instances where flies only momentarily extended or did not extend their legs.

### Head saccade detection

To detect flight saccades, we used an Angular-Velocity based Saccade Detector (AVSD), which works by analyzing the angular heading of the body/head (Censi et al., 2013). We implemented an AVSD with parameters tuned specifically to *Drosophila* head movements (Figure Supplement 1). For each trial, we computed the velocity of the head using the central difference formula (Figure Supplement 1A) and then applied a 10 Hz low-pass filter to the velocity signal (Figure Supplement 1B). We pooled the filtered velocity data points across all trials and computed the standard deviation (Figure Supplement 1G). For each trial, we selected a saccade threshold equal to 1.5 of the overall standard deviations from the trial’s median velocity. For experiments with a static background and sinusoidal stimuli, we used the median of the absolute value of the velocity signal. We used the MATLAB *findpeaks* routine to detect spikes in the filtered velocity signal which were considered putative saccades (Figure Supplement 1B). We also ensured that all spikes lasted within the expected time frame for saccades (10–100 ms). Because we detected peaks in the filtered signal and the spike dynamics were therefore attenuated, we set a window around each peak in the raw signal to find the true peak (Figure Supplement 1C). We applied another threshold set at 2 standard deviations from the median of the raw velocity at this step to ensure that we did not mistakenly detect the faster smooth intervals (such as for 150°s^−1^ stimuli) (Figure Supplement 1G). We determined the start and end of a saccade by finding when the velocity returned to 25% of the peak velocity (Figure Supplement 1C). Saccades where the head moved less than 4° were rejected to avoid detecting the occasional large velocity spike during smooth intervals. Saccades less than the amplitude and/or the velocity threshold were exceedingly rare as shown by the distribution of saccade dynamics (Figure 2B). Furthermore, the velocity and amplitude thresholds were visually confirmed to be effective on ~1000 experimental trials. Next, we extracted the position and velocity of the head during each saccade and interval-saccade interval (smooth movement) (Figure Supplement 1D). We normalized saccades to their peak time and interval-saccade intervals to their start time for later analysis (Figure Supplement 1E). Intervals at the start and end of trials were not bounded by saccades and were therefore not included in our analysis. We removed saccades by shifting each smooth interval by the negative value of the previous saccade amplitude and interpolating between subsequent intervals (Figure Supplement 1F). This procedure allowed us to approximate the smooth dynamics of the head (Figure 9B).

### Wing saccade detection

Prior work has demonstrated that flies attempt to perform body saccades while rigidly tethered (Wolf and Heisenberg, 1980). These fictive saccades manifest as torque spikes which are produced by hitches in ΔWBA. We refer to these hitches as wing saccades. While head saccades are relatively straightforward to detect, wing saccades were much less discernible due to the larger measurement noise floor. This was exacerbated for trials with a high saccade rate (e.g. Video 4). We found that manually detecting wing saccades was not feasible, as wing saccades were not nearly as visually distinct as head saccades overall. Prior studies have attempted to classify wing saccades using algorithms that focus on the entire above-baseline spike in ΔWBA (Aptekar et al., 2012; Bender and Dickinson, 2006), however here we show that head and wings saccades may be best classified by analyzing the *change* in ΔWBA (Figure 7). We found that we could use the same angular velocity-based algorithm we applied for head saccade for wing saccade detection (Figure Supplement 1). We low-pass filtered wing data with an 8 Hz cut-off frequency and set the amplitude cut-off to 4°. We only culled wing saccades for trials with 30°s^−1^ stimulus velocity, as the lower saccade rate and noise floor facilitated detection.

### Protocol to measure passive response

To measure the passive response of the head, we anesthetized flies using the anesthetic agent triethylamine (commercially available as FlyNap, Carolina Biological Supply) which immobilized flies for approximately 45 minutes. We mounted the anesthetized flies on a rigid rod by applying a small drop of UV-activated glue (XUVG-1, Newall) to the thorax, similar to the experiments described above. We then displaced the head using a rod oriented to one side of the fly (Figure 3B). We displaced the head using a manual micrometer while observing displacements with a camera which provided a high-resolution view of the head-neck joint. We ensured that the majority of the displacement was due to the angular offset of the head, as opposed to translational offsets at the base of the neck. The initial angular displacement was varied from realistic values at the onset of head reset saccades (Figure 2C; less than 20°) to displacements that are less naturalistic (range: 20°–40°). However, the initial displacement had little effect on our conclusions, specifically the estimate of the time constant (Figure Supplement 3). After displacing the head, we rapidly moved the rod holding the head away from the fly, allowing the passive response to begin. We recorded videos of the fly from above at 300 frames per second with an infrared-sensitive camera (acA640-120gm, Basler) (Video 3). Each head response was measured using custom computer-vision software (described above). A first-order exponential model of the form

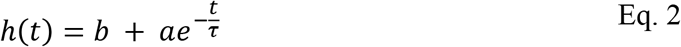

was fit to the experimental data (Figure 3). The head response *h*(*t*) was modeled as a function of time *t*, and free parameters *a, b*, and time constant *τ*. The coefficient of determination (R^2^) fit was greater than 0.9. The active saccade response was fit with the same model form and had an R^2^ in the same range as the passive response.

### Statistical Analysis

For all box plots, the central line is the median, the bottom and top edges of the box are the 25th and 75th percentiles and the whiskers extend to ±2 standard deviations. Violin plots (Figure 4D) used a kernel density function to approximate the distribution. When performing linear regression, we report the Pearson correlation coefficient (*r*). The sigmoid fit to the cumulative distribution function (Figure 6A) is of the form

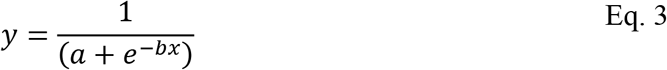

where *a* and *b* are free parameters.

## Acknowledgments

We thank Susheel Dharmadhikari for laboratory assistance. This material is based upon work supported by the Air Force Office of Scientific Research under award number FA9550-20-1-0084.

## Author contributions

B.C. and J.M.M. contributed to the design of experiments. B.C. performed the experiments. B.C. and W.S. analyzed the data. B.C. and J.M.M. wrote the manuscript.

## Supplementary Materials

### Supplementary Figures

**Figure Supplement 1.**
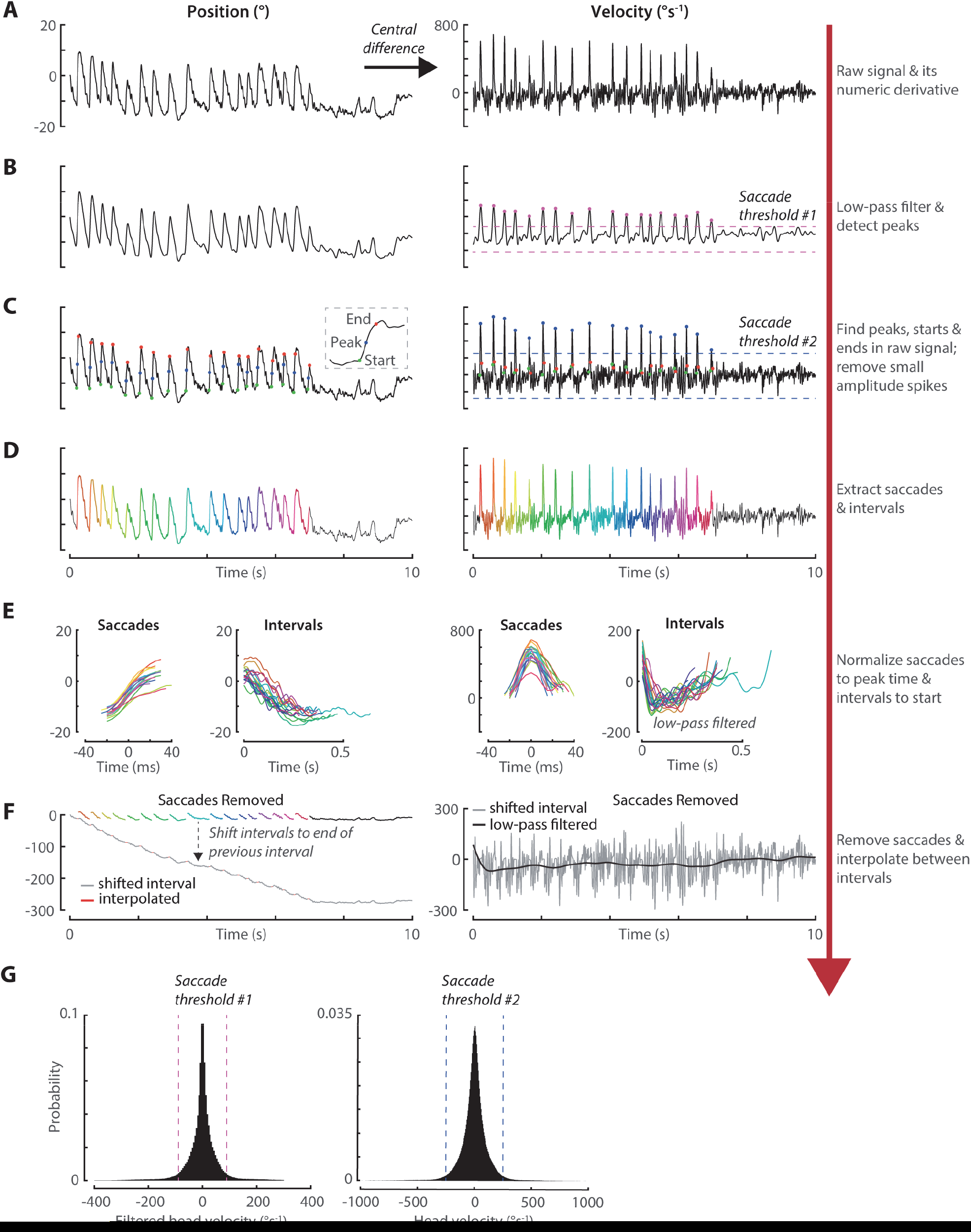
Head saccade detection method. **A)** Raw head trajectory in response to a 150 °s^−1^ counterclockwise rotating background at 30° spatial wavelength (left). We estimated the head’s velocity using the central difference formula (right). **B)** Low-pass filtered head response. We detected peaks in the head’s filtered velocity signal using a thresholding algorithm. Pink markers indicate the peak velocity of putative saccades and the dashed lines represent the detection thresholds in both directions. **C)** The raw head response with the true peaks indicated by blue markers. In order to be considered saccades, velocity spikes had to pass a second threshold indicated by the blue dashed lines. Green and red markers indicate the start and end of each saccade respectively. **D)** The raw head response with all saccades highlighted in distinct colors and the preceding smooth inter-saccade interval highlighted in a dulled version for the same color. The black line indicates data that is neither a saccade nor an inter-saccade interval. **E)** Extracted saccades and inter-saccade intervals using the same color scheme as D). Saccade were aligned to their peak time and intervals aligned to their start time. **F)** The raw head response with saccades removed and each inter-saccade interval shifted to the head position right before the succeeding saccade was triggered (left). The grey curve indicates the shifted intervals while the red curve indicates the interpolated data the replaces each saccade. The original saccades are shown with the same color scheme as D). The black line overlayed on the raw head velocity (right) is the low-pass filtered response, demonstrating a quasi-continuous velocity when saccades were removed. **G)** The distribution of head velocity data points (left) and filtered velocity data points (right) with the saccade detection threshold indicates by the dashed lines with the same color scheme as B) and C).

**Figure Supplement 2.**
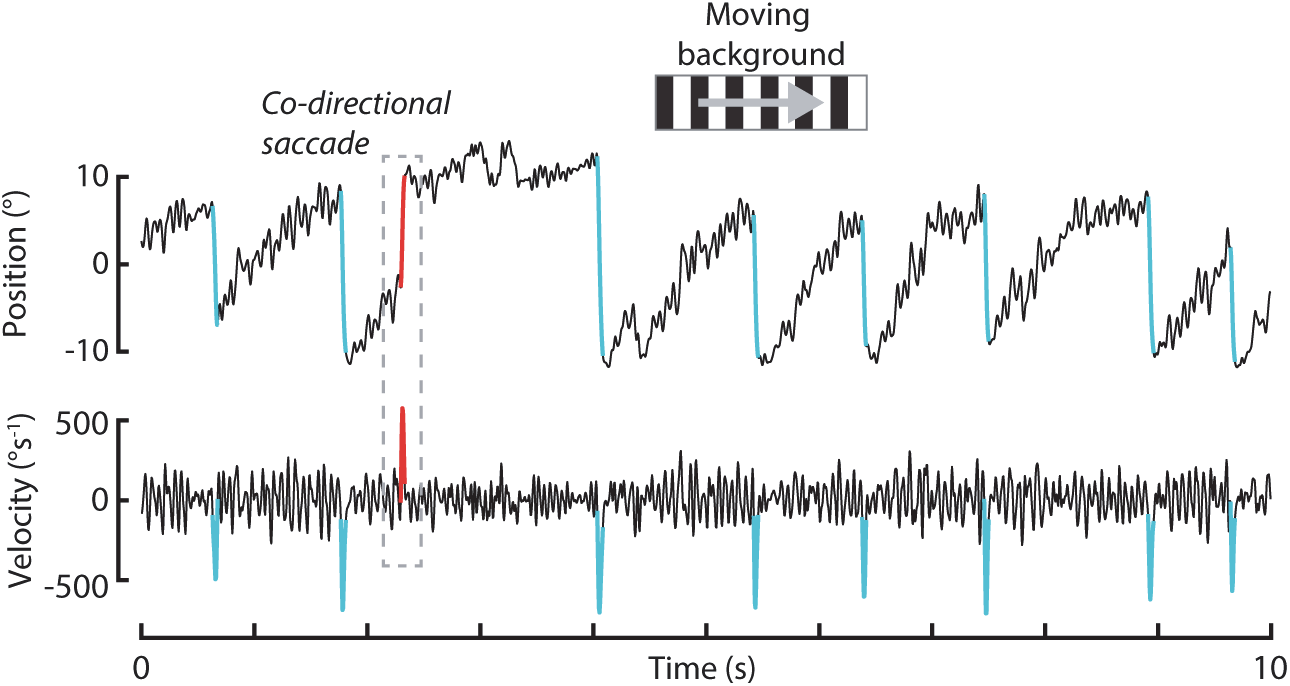
Co-directional saccade. Example head trajectory in response to a 30 °s^−1^ clockwise rotating background at 30° spatial wavelength. Anti-directional (reset) saccades are highlighted in cyan while the one co-directional saccade is in highlighted red. Co-directional saccades were rarely observed (making up less than 2% of a saccades) and therefore this trial is not representative of our data.

**Figure Supplement 3.**
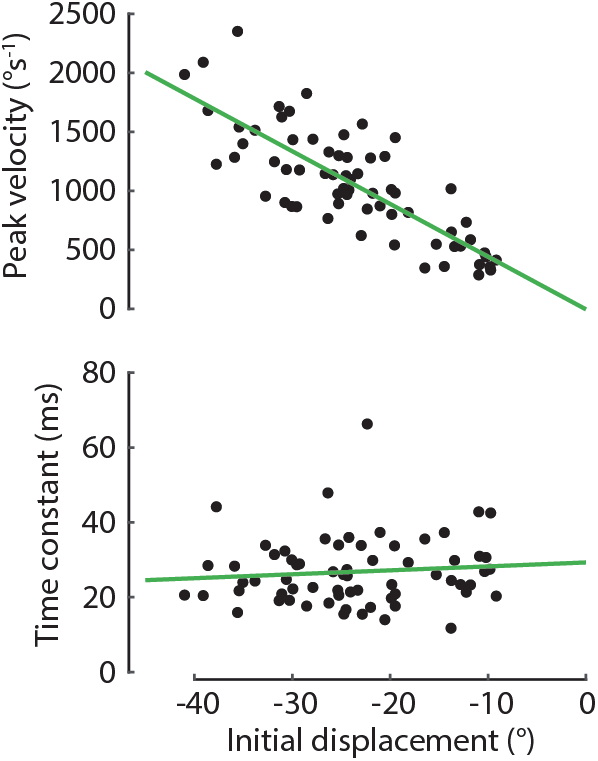
Passive mechanics response, related to Figure 3. The peak velocity (top, *p* < 0.001) and fit, first-order order time constant (bottom,*p* = 0.4) of the head’s free response for various initial displacements. Green lines indicate the linear best fit. *n* = 6 flies.

**Figure Supplement 4.**
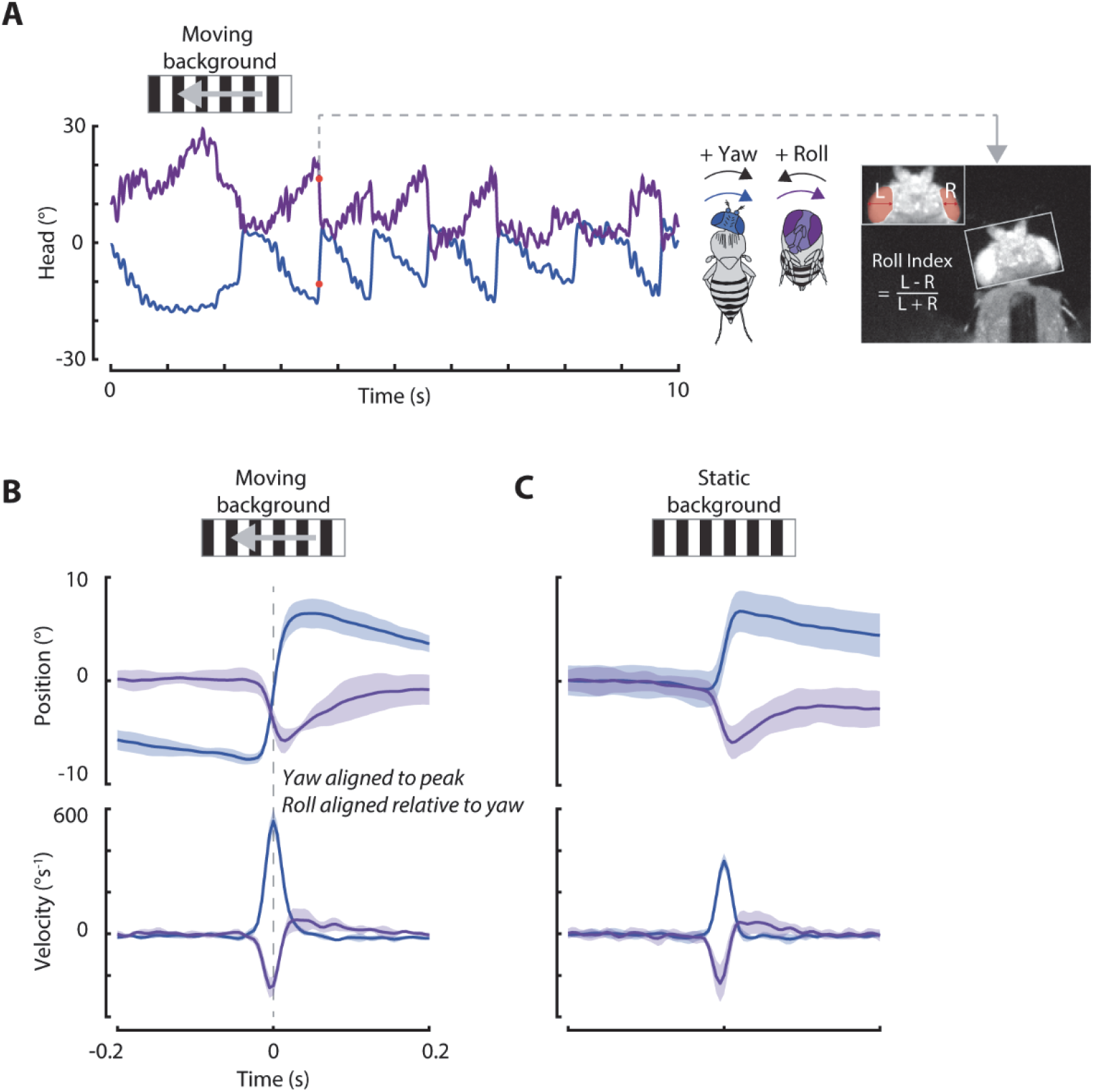
Head roll during saccades. **A)** Example head trajectory in response to a 30 °s^−1^ counterclockwise rotating background at 30° spatial wavelength. Head yaw (blue) smoothly follows the stimulus and saccades move in the opposite direction. During each clockwise yaw saccade, the head rolls (purple) counterclockwise. The roll index was estimated using the ratio of the projected widths of each eye. Inset: example of the head at a large yaw and roll orientation (dorsal view). Same experimental trial as shown in Video 2. **B)** The head yaw position and velocity (blue) aligned to head yaw saccade velocity peak time, and head roll position and velocity (purple) aligned relative to yaw, such that the true time difference between yaw and roll was maintained. Stimulus: 30°s^−1^ rotating background at 30° spatial wavelength. Counterclockwise saccades were inverted and pooled with clockwise saccades. **C)** Same as B) but for saccades detected during the presentation of a static background. Shaded regions: ±1 standard deviation. 30°s^−1^ background: *n* = 5 flies, *N* = 809 saccades. Static background: *n* = 8 flies, *N* = 547 saccades.

**Figure Supplement 5.**
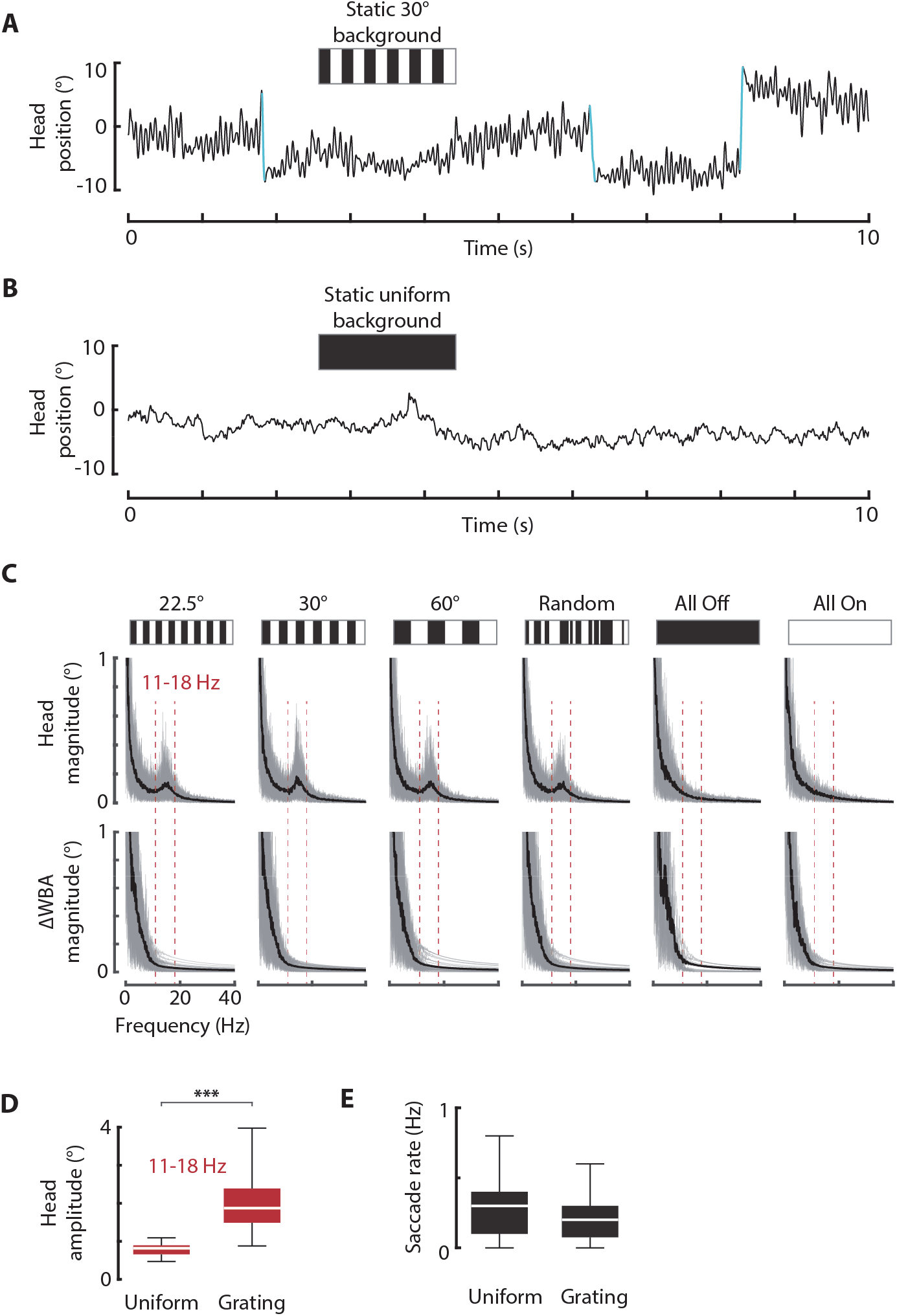
Spontaneous head movements. **A)** Example head response for a static background with 30° spatial wavelength. Note the small oscillatory head movements between saccades. Saccades are highlighted in cyan. **B)** Same as A) but for a uniform (dark) background. Note the absence of oscillatory head movements. **C)** The frequency domain representation of head (top) and wing (bottom) movements (calculated via a Fast Fourier transform) for flies presented with gratings of various spatial wavelengths. Grey lines: individual trials. Black curve: mean response. Shaded area: ± 1 STD. Note the difference in the head response for gratings with salient visual features and uniform gratings between 11–18 Hz (dashed red lines). **D)** The distribution of the amplitude of oscillatory head movements between 11–18 Hz for gratings with salient features (left) and uniform gratings (right). *** = *p* < 0.001. **E)** The distribution of the head saccades rates for gratings with salient features (left) and uniform gratings (right). *n* = 8 flies.

**Figure Supplement 6.**
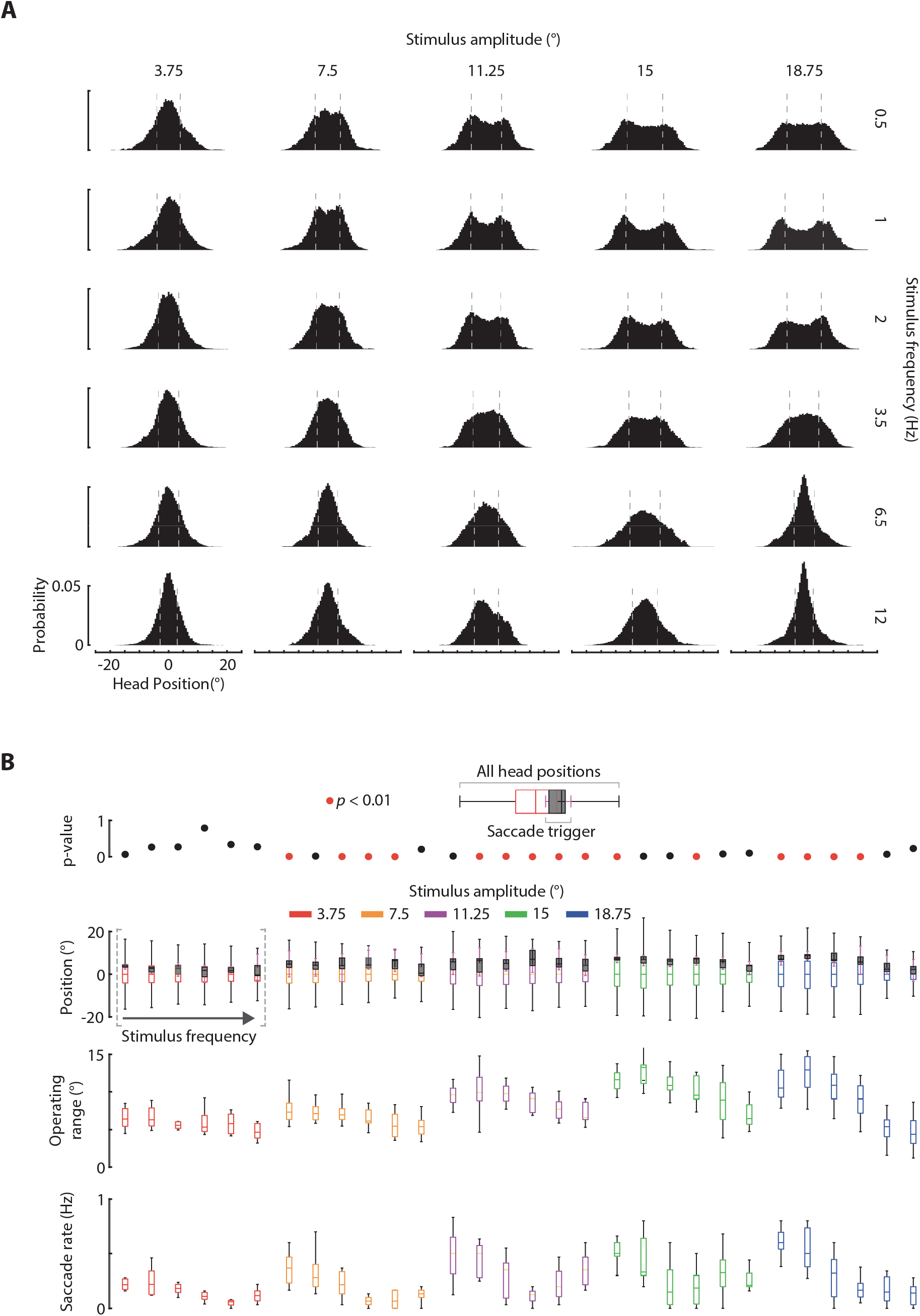
Head operating range, related to Figure 6. **A)** The distribution of all head orientations for sinusoidal visual stimulus at distinct amplitude-frequency combinations. The dashed grey lines indicate the operating range of the head for each distribution, calculated as twice the mean of the absolute value of all data points. **B)** The total head orientation distributions (colored) compared to the distributions of saccade start orientations (black, normalized for counterclockwise saccades). The *p*-values comparing the total head distribution to the saccade start orientations are shown above and the operating range and saccade rate distributions are shown below. 3.75°: *n* = 8 flies, *N* = 746 saccades. 7.5°: *n* = 10 flies, *N* = 602 saccades. 11.25°: *n* = 11 flies, *N* = 957 saccades. 15°: *n* = 10 flies, *N* = 1,068 saccades. 18.25°: *n* = 11 flies, *N* = 1,057 saccades.

**Figure Supplement 7.**
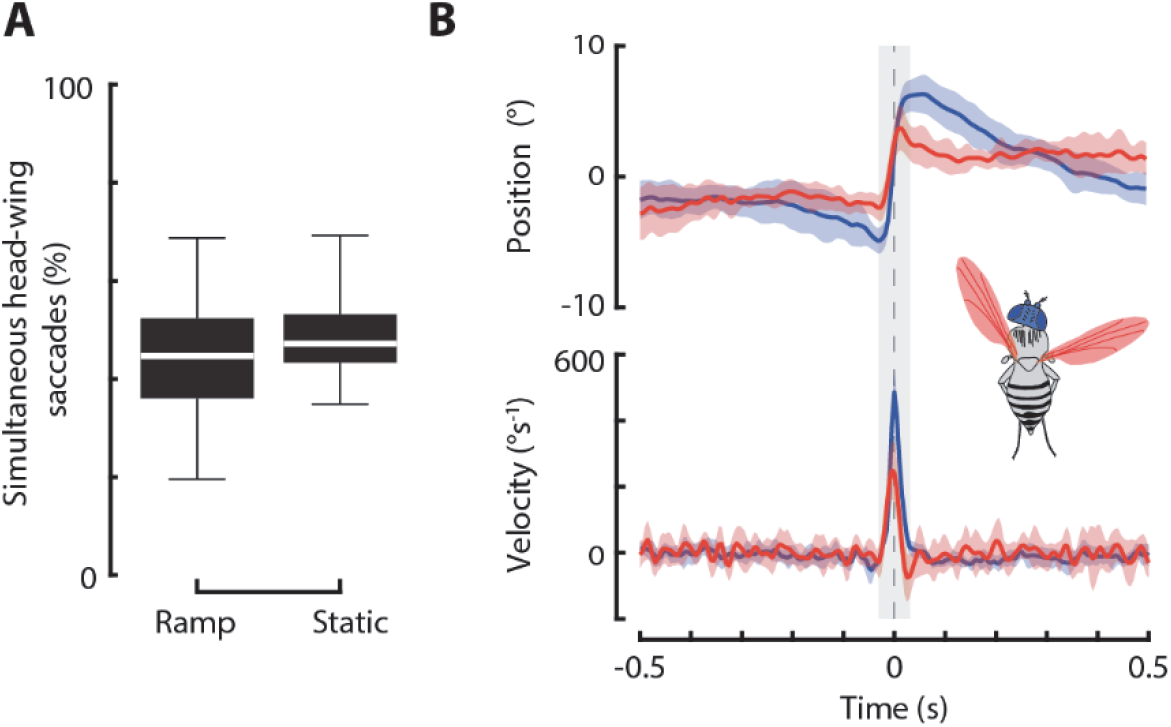
Head and wing saccades, related to Figure 7. **A)** The distribution of the percent of head saccades that had an identifiable wing saccade within 100 ms for ramp visual stimulus and for a static background. Ramp: *n* = 12 flies, *N* = 829 head saccades. Static background: *n* = 8 flies, *N* = 359 saccades. **B)** The head and wing response for head saccades that did not have an identifiable wing saccade within 100 ms for ramp visual stimulus. Note the smaller amplitude of the wing response, which made detecting these potential saccades highly challenging without a correspondingly higher false positive rate. Counterclockwise saccades were inverted and pooled with clockwise saccades. Shaded regions: ±1 standard deviation. *n* = 12 flies, *N* = 302 head saccades.

**Figure Supplement 8.**
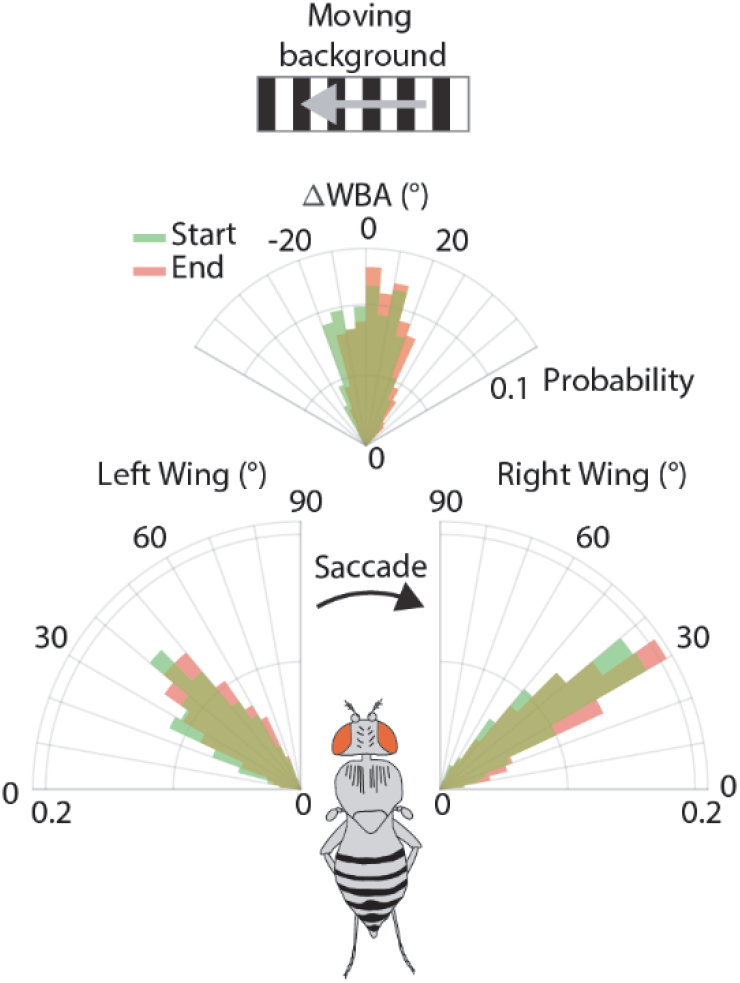
Wing orientation at saccade initiation, related to Figure 8. The distribution of left and right wingbeat amplitudes (left & right respectively; bottom) and ΔWBA (top) at the start (green) and end (red) of each head saccade. Counter-clockwise saccade ΔWBA were inverted and pooled with clockwise saccade ΔWBA. Similarly, left- and right-wing WBA were normalized so that all wing saccades were clockwise. Compared to head start and end distributions (Figure 2C), there is a significant overlap between the distributions across left, right, and ΔWBA data. *n* = 12 flies, *N* = 829 head saccades.

### Movie Captions

**Video 1.** A real-time experimental trial showing head and wing movements in response to constant velocity visual motion at −60°s^−1^ and 30° spatial wavelength (data same as Figure 1C). The captured video of the fly is shown inside an artificially recreated virtual reality arena (not to scale) with head and wing tracking overlaid on top of the recorded video. The corresponding motion of the visual display, head, and ΔWBA are shown to the right, synchronized with the video. The video was captured at 200 frames per second but subsampled to 50 frames per second for display purposes.

**Video 2.** Illustration of our methods of estimating roll from a top view of a fly.

**Video 3.** Passive mechanics experiment. The head is displaced and released, and the corresponding free response is shown. Shown three times in real time and once slowed down by ~6X.

**Video 4.** Same as Video 1 but for 150°s^−1^ visual motion at 22.5° spatial wavelength.

**Video 5.** Same as Video 1 but for a static background at 30° spatial wavelength.

**Video 6.** Same as Video 1 but for no spatial features in near-complete darkness (LED lights off).

**Video 7.** Same as Video 1 but for 150°s^−1^ visual motion. Note the extension of the fly’s legs tracked by markers on the left and right front legs.

**Video 8.** Same as Video 1 but for 120°s^−1^ visual motion. Note the switch from stabilizing head movements and saccades to a landing response.

**Video 9.** Same as Video 1 but for sinewave visual motion at 3.75° amplitude and 2 Hz oscillation frequency. The stimulus position is also shown on the same axes as head position, indicated by the green curve.

**Video 10.** Same as Video 1 but for 18.75° amplitude.

**Video 11.** Same as Video 1 but for 6.5 Hz oscillation frequency.

**Video 12.** Same as Video 1 but for 12 Hz oscillation frequency.

**Video 13.** Top: same as Video 1 but for 30°s^−1^ visual motion. Head and wings are plotted on the same axes for comparison. Bottom: Same as top, but for a head-fixed fly.

